# Mitotic Activity and DNA Maintenance of Adult Neural Stem Cells is Regulated by Beclin1

**DOI:** 10.1101/2024.01.26.573092

**Authors:** A. Kalinina, J. Dhaliwal, Y Xue, M. Vaculik, M. McCambley, B.C. Fong, D. Cook, R.S. Slack, D.C. Lagace

## Abstract

Beclin1 is a tumor suppressor gene that can regulate proliferation under pathological conditions. Whether Beclin1 has a role in regulating proliferation in physiological conditions remains unknown. Here, through the creation of an inducible transgenic mouse that removes Beclin1 from adult neural stem and progenitor cells (NSPCs) and their progeny we uncovered that Beclin1 is required cell-autonomously to sustain proliferating NSCPs *in vivo* and *ex vivo*. Flow cytometry analysis and single-cell RNA-sequencing show that Beclin1 is required for mitosis. Additional analysis of the proliferating NSPCs resolved by stage of cell cycle uncovered that Beclin1-null cells have a distinct differential developmental trajectory accompanied by downregulation of genes involved in chromosomal maintenance and upregulation of cell stress genes upon cell cycle exit. These effects align with DNA damage in Beclin1-null cells and ultimately result in less adult-born granular neurons. Together these data identify Beclin1 as a novel regulator of mitosis in adult NSPCs.

## INTRODUCTION

Beclin1 (Atg6) is a scaffold protein that has well-established roles in the regulation of autophagy^1,2^, endocytosis^3,4^, and phagocytosis^5,6^. There is also a growing appreciation that Beclin1 is multifunctional and can regulate proliferation within the field of oncology, where it has been characterized as a tumor suppressor. For example, in aggressive breast cancer cells, knockdown of Beclin1 suppresses proliferation by regulating the cell cycle and apoptosis^7^. Similarly, in colorectal and HeLa cells Beclin1 deficiency disrupts chromatin dynamics during mitosis resulting in deficient proliferation^10^, and increases genomic instability^8–10^. Whether Beclin1 regulates proliferation under normal physiological conditions remains largely unknown.

The examination of the role in Beclin1 has been mainly studied in the embryonic and early stages of development. Homozygous Beclin1 knockout mice are embryonic lethal in early stages of development^11^. Heterozygous Beclin1 knockout mice are viable but have significant increases in tumorgenicity^11^ as well as a reduction in neurogenesis in the olfactory bulb^12^. Conditional Beclin1 mouse models have further shown that loss of Beclin1 in keratinocytes is lethal due to dampened expansion of the embryonic epidermis^4^, while loss of Beclin1 in select neurons during development leads to severe neurodegeneration and premature death^13^. Together this body of literature suggests that Beclin1 may be necessary for proliferation *in vivo* under physiological conditions. To test this hypothesis, we specifically explore the requirement of Beclin1 within the neural stem and progenitor cells (NSPCs) that develop into adult granule neurons within the hippocampus of the adult brain.

Our findings reveal that Beclin1 is required during adult neurogenesis through its role in mitosis and chromatin stability. Through the creation of an inducible Beclin1 knockout mouse, Beclin1 removal from adult nestin-expressing NSPCs and their progeny resulted in a significant cell-autonomous reduction in proliferating NSPCs. The combination of flow cytometry and single-cell RNA sequencing (scRNA-seq) analysis independently highlighted Beclin1 was required during mitosis. The ability to resolve the states of proliferating NSPCs, as well as cells exiting the cell cycle identified that Beclin1-null mitotic cells have a distinct differential devel-opmental trajectory accompanied by changes in key players that regulate mitosis and differentiation, and directly modify genes impeding proliferation. Specifically, mitotic cells downregulate genes involved in chromosomal maintenance and upregulate cell stress genes upon cell cycle exit. These deficits are accompanied by DNA damage in Beclin1-null cells which ultimately decreased their survival and led to a reduction in the generation of adult-born neurons.

## RESULTS

### Loss of Beclin1 reduces proliferation cell-autonomously

To conditionally remove Beclin1 from adult nestin-expressing NSPCs and progeny, we created a Beclin1 Nestin-inducible transgenic mouse model (hereafter referred to as Beclin1 nKO; NestinCreER^T2^:R26R-eYFP: Beclin1^flox/flox^) and littermate control wild-type (WT) mice (NestinCreER^T2^:R26R -eYFP: fBeclin1^WT/WT^). The recombined (YFP+) cells in the Beclin1 nKO mice were confirmed to have a significant reduction in the amount of Beclin1, as shown by the quantification of Beclin1 puncta in fixed brain sections (Fig 1A,B), as well as western blot detection of Beclin1 protein from YFP cells purified using fluorescence-activated cell sorting (FACS) (Fig 1C). To further test if the Beclin1-null cells had a reduction in autophagy, Beclin1 nKO and WT mice were injected with a Cherry-EGFP-LC3 retrovirus, using our published *in vivo* methodology^14^. Beclin1-null cells in the Beclin1 nKO mice had a significant reduction in autolysosomes (mCherry+ puncta) compared to control cells in the WT mice (Fig S1 A,B), supporting the role of Beclin1 in regulating autophagy^1,2^.

**Figure 1.**
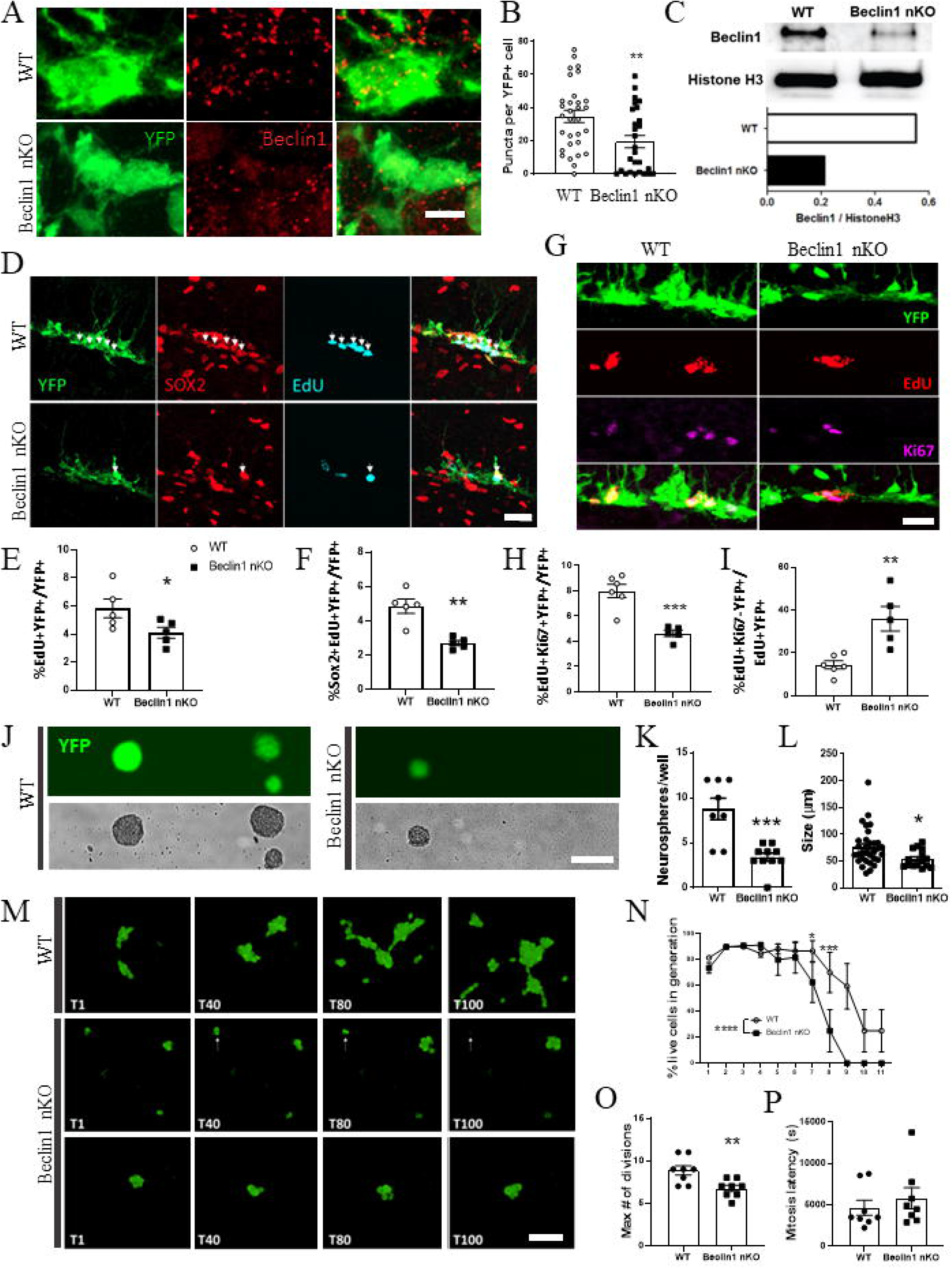
Removal of Beclin1 from adult NSPCs reduces proliferation *in vivo* and *ex vivo*. (A) Representative images and (B) quantification of Beclin1 protein puncta (red) in YFP+ cells (green) showing a reduction in puncta in Beclin1 nKO mice. (C) Western blot analysis of Beclin1 protein, normalized by Histone H3 in pooled YFP+ cells, showing less Beclin1 protein in Beclin1 nKO mice. (D) Representative images and quantification of YFP+ (green) cells showing a reduction in (E) proliferating NSPCs (EdU+, blue), as well as proliferating EdU+ Sox2+ (red) NSCs in Beclin1 nKO mice. (G) Representative images and quantification (H, I) of YFP+ cells labeled with EdU (red) and Ki67 (violet) showing that of the total number of all YFP+ NSCPs there is a reduction in the (H) activity of cycling (EdU+,Ki67+) cells and (I) increase in cells exiting the cell cycle (EdU+Ki67-) in the Beclin1 nKO mice. (J) Representative images of YFP+ neurospheres (green, top) and brightfield images (bottom) of primary cultures made from WT and Beclin1 nKO mice showing a significant reduction (K) of total numbers of neurospheres per well, and (L) neurosphere size. (M) Representative images from live imaging showing cell division over time (T1-T100) with WT cells growing *in vitro* exponentially, while Beclin1 nKO cells undergoing cell death (white arrow, middle panel) and reduced expansion (lower panel). (N) Percentages of live cells generated via cell division across 11 generations in WT and Beclin1 nKO samples showing a significant drop in generation of cells in Beclin1 nKO samples after generation 6. (O) Beclin1 removal reduced the maximum number of NSPC divisions in the absence of (P) altering mitotic latency. Scale bars represent 5 uM (A), 20 uM (D), 20 uM (G), 100 uM (J), 100 uM (M); For all experiments mice were harvested at 14 dpi with the exception of C (harvested at 35 dpi); All graphed data show individual animals (B, E,F,H, I), or wells/neurospheres/cells (K, L, O, P) as well as the mean ± SEM, with the exception of pooled cells in C; data with 2 groups were analyzed by unpaired t-test, with exception of (N) analyzed by 2-way repeated ANOVA; * p≤0.05, ** p≤0.01, *** p≤0.001, **** p≤0.0001.

We next assessed proliferation in the Beclin1 nKO compared to WT mice. To label the slowly dividing activated NSCs (aNSCs) and rapidly dividing NPCs^15–18^, two weeks after the injection of tamoxifen (14 dpi), the mice were treated with multiple injections of EdU (5-ethynyl-2′-deoxyuridine) two hours apart and perfused two hours after the last injection^19^. EdU incorporation showed a significant genotype-dependent decrease in the proportion of proliferating (YFP+Edu+) Beclin1-null cells (Fig 1D, E). Additionally, the proportion of aNSCs, identified by their expression of EdU and a stem cell marker, Sox2^20–22^, showed a significant decrease in Beclin1-null aNSCs (Fig 1D, F). This decrease in the proportion of aNSCs was not associated with an increase in the proportion of Beclin1-null quiescent NSCs (qNSCs), as there were no significant differences in the proportion of qNSCs between the Beclin1 and WT mice (Fig S2A, B). These findings suggest that Beclin1 reduces the proliferation of aNSCs and NPCs.

To examine if Beclin1 is required in aNSCs and NPCs that are actively cycling or exiting cell cycle, Beclin1 nKO and WT mice were treated with multiple injections of EdU two hours apart and were perfused 24 hours after the first injection^19^. Beclin1 nKO mice had a significant reduction in the proportion of cells that were actively cycling as measured by the YFP+EdU+ cells that expressed Ki67 (Fig 1G, H). Additionally, Beclin1 nKO mice had a significant increase in the proportion of EdU+ cells exiting the cell cycle (EdU+Ki67-), with an average of 36% of EdU-labeled Beclin1-null cells leaving cell cycle in a 24-hour period compared to only 14% of WT cells (Fig 1G, I). Together these data support the notion that loss of Beclin1 reduces proliferation in the actively cycling aNSCs and NPCs.

To begin to determine if the reduction in proliferation in the Beclin1-null cells was due to a cell-autonomous mechanism the *in vitro* neurosphere assay was performed using cultures obtained from the dentate gyrus of adult Beclin1 nKO and WT adult mice. Both the number and size of the primary YFP+ spheres were significantly reduced in the Beclin1 nKO compared to WT mice (Fig 1J-L). To further examine the dynamics of cell division of daughter cells in Beclin1-null cells we performed live cell imaging of WT and Beclin1-null DG cells in monolayer culture conditions. WT cells created healthy progeny as observed by their maintained rate of proliferation throughout the imaging period producing 11 generations (Fig 1M, N). In contrast, Beclin1-null cells showed increased cell death over time and did not produce daughter cells after 8 generations. This was accompanied by a significant 25% reduction in the maximum number of divisions that could be carried out by Beclin1-null compared to WT cells (Fig 1O). In addition, WT and Beclin1-null cells that successfully underwent mitosis were not different in latency to enter next mitosis (Fig 1P) suggesting that Beclin1 removal does not perturb cell abscission or entry into next cell cycle. Overall, these data support Beclin1 is required in the regulation of NSPC proliferation.

### ScRNAseq analysis highlights altered mitotic dynamics of Beclin1-nell cells

To gain insights into the cellular mechanisms underlying the disrupted proliferation and increased cell death in the Beclin1 nKO mice, we dissected and collected single YFP+ WT and Beclin1-null cells by FACS and subjected them to the 10x Genomics single cell RNA sequencing (scRNAseq). As expected, examination of the WT and Beclin1-null cells revealed that the lineage of recombined nestin-expressing neurogenic cells and their progeny formed the largest group of related cells (Fig 2A). In addition to these cells, three other small groups of cells that are known to express Nestin were detected, including pericytes^23^, endothelial cells (EC)^23^, and myelin-forming oligodendrocyte (MFOL) precursor cells^24^.

**Figure 2.**
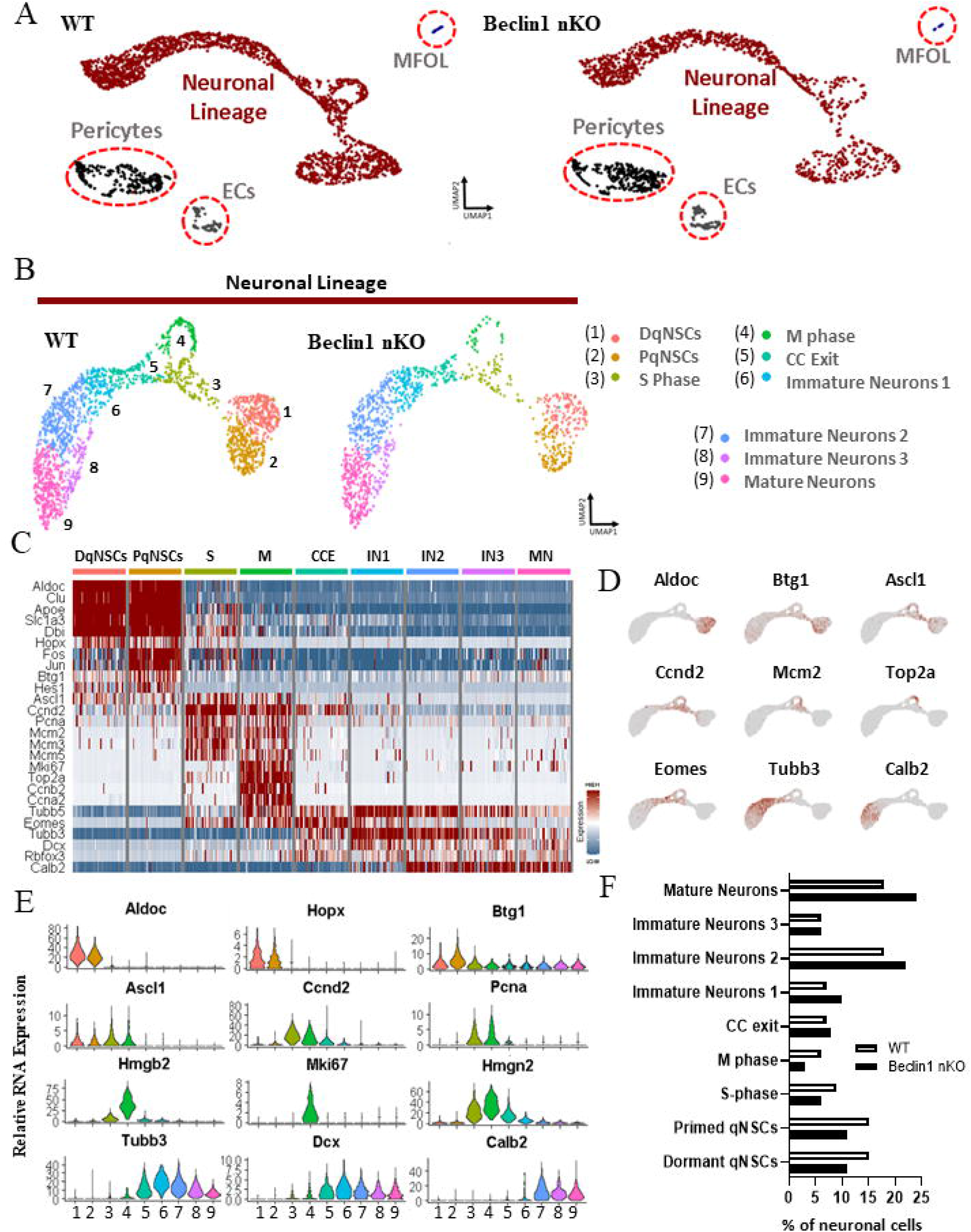
sc-RNAseq of WT and Beclin1 nKO mice highlights heterogeneous neurogenic populations. (A) UMAP projection of WT and Beclin1 nKO sc-RNAseq samples generated from cells sorted at 14 dpi. (B) UMAP projection of the neuronal lineages without other cell types reveal clusters of dormant quiescent stem cells (DqNSCs), primed qNSCs (PqNSCs), S phase cells, M phase cells, cells exiting cell cycle (CC Exit/CCE), immature neurons 1-3, and mature neurons. (C) Heatmap of relative RNA expression of cluster markers within the neurogenic lineage shows high specificity of grouping by cell type and state (blue=low, red=high). (D) Examples of cell markers’ expression shown in UMAP projections. (E) Violin plots of relative RNA expression of known markers used to identify different cell types showing Aldoc, Hopx, and Btg1 expression in dormant and primed qNSCs, Ascl1, Ccnd2, Pcna, Hmgb1, Mki67, Hmgn2 expression in proliferating stem and progenitor cells, and Tubb3, Dcx, and Calb2 in immature and mature neurons. (F) Percentages of different cell clusters within the neuronal lineage of WT and Beclin1 nKO samples shows a decline in mitotic cells.

Analysis of the neuronal lineage of the WT and Beclin1 nKO samples allowed for the resolution of 9 different cell groups in the neurogenic lineage (Fig 2B). The qNSCs were detected by Aldoc, Hopx, and other quiescence genes^25^ (Fig 2C-E). They were divided into dormant qNSCs (DqNSCs) and primed qNSCs (PqNSCs), with the PqNSCs having more expression of genes such as Btg1 and Fos^26^. Using the transient expression of cell cycle markers, the proliferating NPCs and NSCs were divided into cells in S phase and G2/M phases (hereafter, M phase) of the cell cycle (Fig 2C). S phase cells were marked by genes such as Ascl1, Mcm2, and cyclin D2 (Ccnd2), while M phase cells were positive for Top2a, Hmgb2/n2, and Mki67 (Fig 2C-E). There was also a group of cells exiting cell cycle (CCE) which were marked by a large reduction in Pcna^27^ and rise of early differentiation markers (Fig 2C-E). The remaining non-cycling neurogenic cells consisted of four clusters of neurons. They included three groups of immature neurons labeled IN1-3 that had decreasing expression of immature neuron markers Eomes, Tubb3, and Dcx (Fig 2C-E). The last group of neurons was labeled as mature neurons based on relative increase in expression of mature neuron markers such as Calb2 (Fig 2C-E). Examination of the proportion of these 9 cell types showed the highest change in the proportion of cells in M phase with a 2-fold reduction in the Beclin1-null mice compared to WT mice (Fig 2F). These results led us to hypothesize that Beclin1-null cells have abnormal cell cycle dynamics that mediated the reduction in proliferation.

### Beclin1 nKO mice have fewer mitotic cells *ex vivo* and *in vitro*

To test if the loss of Beclin1 altered the proportion of cells in the different stages of the cellcycle, we performed an *ex vivo* cell cycle assay^28,29^. Specifically, 14 dpi adult WT and Beclin1 nKO mice were administered four injections of EdU two hours apart, then 24 hours after first injection single cells were obtained from the DG. Cells underwent flow cytometry analysis to measure EdU incorporation and DNA content in order to identify cells in S phase, as well as M phase. This assay revealed a significant decrease in the percentage of total mitotic cells in the DG of Beclin1 nKO mice (Fig 3A, B). This reduction in mitotic cells was accompanied by no significant change in the percent of cells in the S phase of the cell cycle (Fig 3A, C). These results therefore verify the scRNAseq results and corroborate the loss of Beclin1-null mitotic cells.

**Figure 3.**
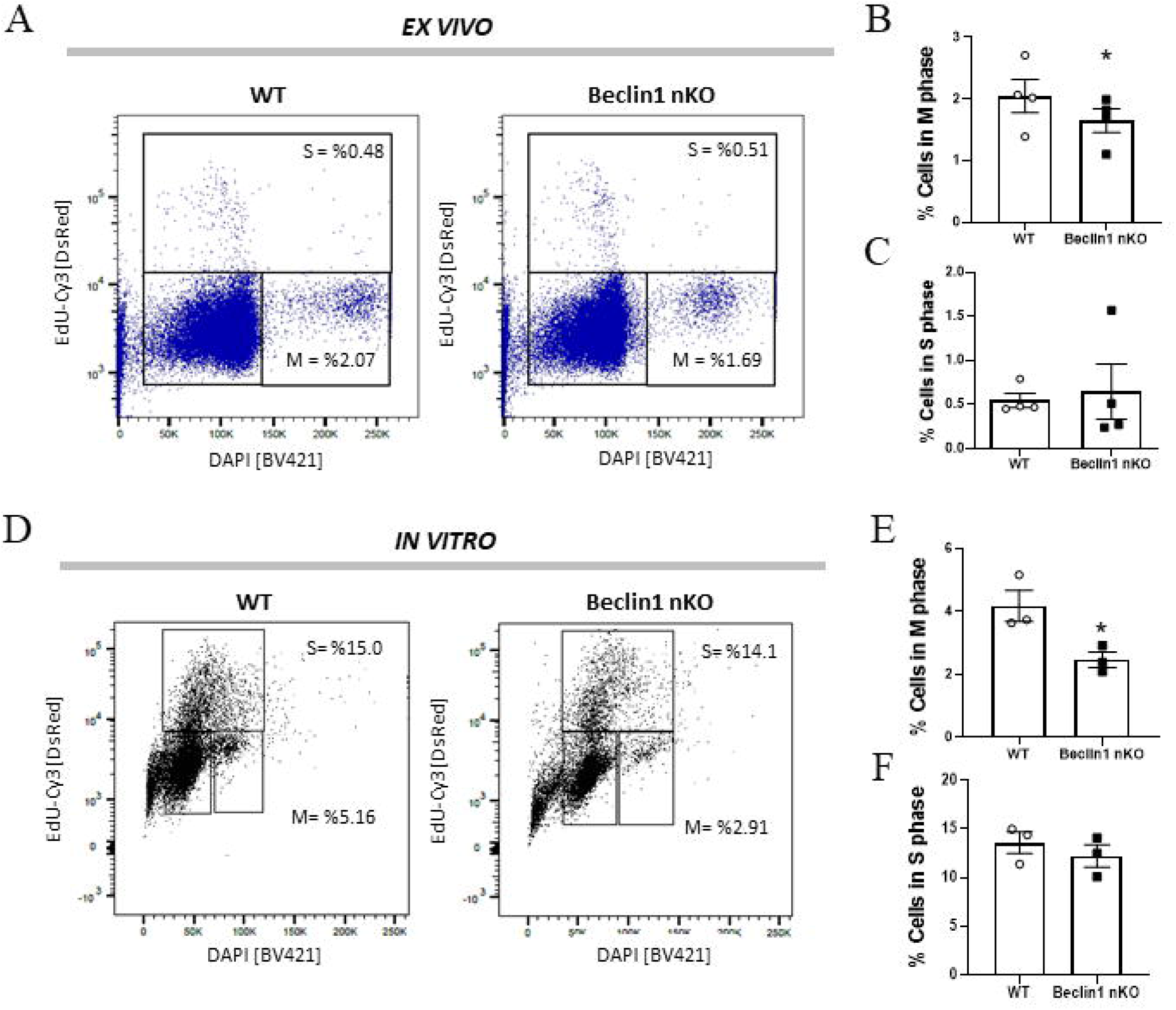
Removal of Beclin1 from adult Nestin NSPCs reduces their mitotic activity *ex vivo* and *in vitro*. (A) Representative images of gating of EdU+ cells in cell cycle flow cytometry graph of EdU-Cy3 (y-axis) and DAPI (x-axis) intensity in cells from WT and Beclin1 nKO mice show (B) a significant reduction of the percentage of cells in M phase in Beclin1 mice, in the absence of change in the (C) percentage of cell in S phase. (D) Representative images of gating of EdU+ cells in cell cycle flow cytometry graph of EdU-Cy3 (y-axis) and DAPI (x-axis) intensity in neurospheres cultured from WT and Beclin1 nKO mice show (E) a significant reduction of the percentage of cells in M phase, in the absence of change in the (F) percentage of cell in S phase. Graphs show relative gene expression in 50 randomly selected cells from each cell cluster (C), relative gene expression values across all individual cells (D), and the distribution of relative gene expression across all cells (E).

To further test if the cell-autonomous reduction in neurosphere formation in Beclin1 nKO mice is accompanied also by a reduction in cells in M phase, we made neurospheres from the DGs of 14 dpi WT and Beclin1 nKO animals and performed an *in vitro* pulse-chase with EdU (Fig 3D). Similar to our *ex vivo* results (Fig 3A-C), the percent of mitotic cells was significantly reduced (Fig 3D, E) in the absence of change in the percent of cells in S-phase (Fig 3F). Together all these data suggest the cell-autonomous reduction in proliferation of Beclin1-null cells is accompanied by altered progression through mitosis.

### Beclin1-null prophase/prometaphase cells are fewer in number and diverge upon cell cycle exit

To explore mechanisms by which loss of Beclin1 impedes mitosis, we resolved the states of proliferating NSPCs identified by our scRNAseq analysis that were in S/M phase or exiting the cell cycle (groups 3-5 in Fig 3B). This revealed eight clusters of cells that could be identified based on their respective stages along the activation-differentiation continuum including: aNSCs; NSCs and NPCs in S phase of the cell cycle; cells in early and late M phase; and 3 groups of cells exiting the cell cycle (Fig 4A, B). To additionally validate the cell cycle status of the proliferating clusters, we used Seurat’s cell cycle scoring algorithm that approximates cell cycle identity as being in G1, G2/M or S-phase based on the expression of known S and M cell cycle genes^30^. The scoring for G1 phase identity aligned with CC1-CC3, whereas the proliferative early/late M cells and S NSCs/NPCs aligned with G2/M and S cells. Additionally, the Seurat cell cycle analysis also validated the reduction in proportion of Beclin1-null cells in the G2/M and S phase (Fig 4A, B, E).

**Figure 4.**
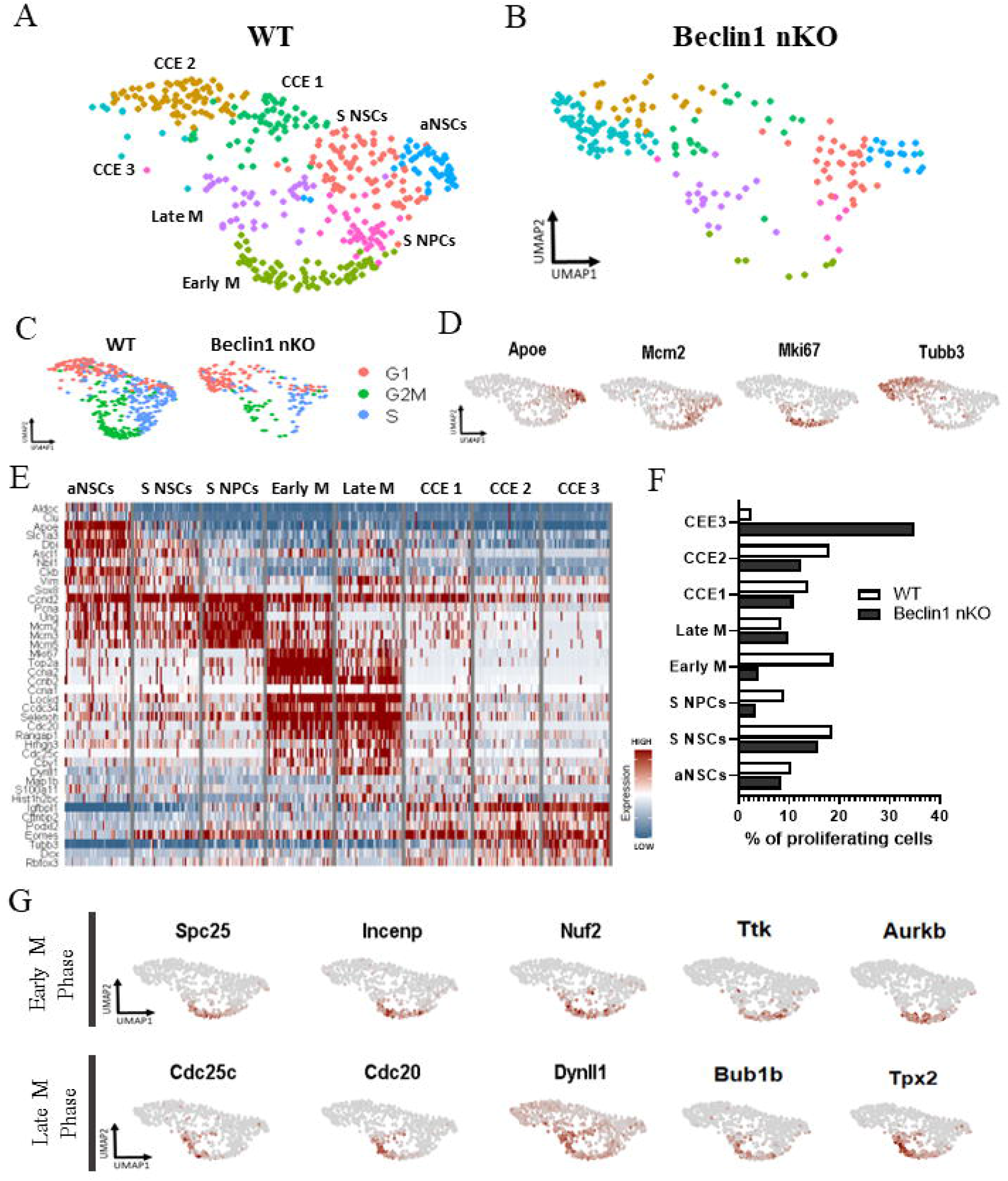
Examination of NSPCs supports differential effect of Beclin1 removal on cell cycle stages. (A, B) Cycling cells (S phase, M phase, and CC exit clusters) were selected for further identification of unique clusters shown in a UMAP projection, including active NSCs (aNSCs) and S phase NSCs (S NSCs), S phase NPCs (S NPCs),G2 and early M phase cells (early M), late M phase (Late M) and cell cycle exit clusters 1-3 (CCE1-3). (C) UMAP projections of the clusters using cell cycle scoring of WT and Beclin1 nKO cells (S in blue, G2M in green, and G1 in red). (D) UMAP projections of marker RNA expressed in cycling cells with Apoe marking aNSCs, Mcm2 marking S phase, Mki67 marking G2M and Tubb3 marking cells exiting cell cycle. (E) Heatmap of relative RNA expression of genes marking the cell types and states (blue=low, red=high). (F) Percentages of different clusters within the WT and Beclin1 nKO samples. (G) UMAP projections of cycling cells from WT and Beclin1 nKO sc-RNAseq samples showing relative RNA expression of prophase/prometaphase (top) and metaphase/anaphase (bottom) genes.

The aNSC subcluster was positive for activation markers Ascl1 and Dbi and NSC markers Aldoc and Apoe (Fig 4D, E). The aNSC cluster was also positive for S phase markers like Mcm genes and Pcna, suggesting the aNSCs represent a transition state between the PqNSCs and S phase NSCs. The aNSCs did not include cells in M phase, as shown by the absence of M phase markers. Between the Beclin1-null and WT cells there was a similar proportion in aNSCs (Fig 4F). The S-phase NSCs had decreased expression of Apoe, Dbi and Ascl1 NSC markers compared to early aNSCs, whereas these genes were completely downregulated in the S-phase NPCs (Fig 4D, E). There was no difference in proportion of S phase NSCs between Beclin1-null and WT cells, yet a prominent decline in the proportion of S phase NPCs (Fig 4F).

Cells in M phase separated into early M and Late M phases, but NSCs and NPCs could not be distinguished from each other (Fig 4A, B). Early M phase cells expressed more prophase, prometaphase, and spindle assembly checkpoint (SAC) genes such as Spc25, Incenp, and Nuf2 (Fig. 4E, G). The Late M cells were expressing SAC, metaphase and anaphase genes like Cdc25c, Cdc20, and Dynll1 (Fig. 4E, G). Compared to WT, Beclin1-null cells had a striking reduction in the proportion of early M phase cells, however, there was no difference in the percentage of cells in late M phase (Fig 4F). This suggests that the cell cycle perturbation in the NSPCs following Beclin1 removal may involve inadequate prophase/prometaphase maintenance.

Lastly, this analysis allowed for the identification of three clusters of cells exiting the cell cycle (Fig 4A, B, CCE 1-3). The differences between CCE1-CCE3 represented early progeny on a continuum of expression of neuronal differentiation markers such as Tubb3 and Dcx (Fig 4D, E). CCE1 has the least amount of neuronal marker expression and likely are the cells that are able to re-enter cell cycle or enter quiescence. In comparison, the CCE2/CCE3 expression high levels of differentiation markers and thus likely differentiate into neurons. The Beclin1-null and WT clustered differently between CCE 1-3. Specifically, Beclin1-null cells preferentially clustered into CCE3, while WT cells localized mostly to CCE2 (Fig 4F), suggesting a differential transcriptional fate for the Beclin1-null compared to WT cells during cell cycle exit.

Together these data highlight the power of scRNAseq analysis to identify robust changes in specific sub-populations of cycling cells that cannot be easily resolved through other methods. For example, our histological analysis of aNSCs showed a reduction in aNSCs (Fig 1D-F), yet the scRNAseq data shows the reduction in aNSCs is due to changes in proportion of aNSCs in M phase, rather than S phase. Additionally, our flow cytometry analysis did not detect a significant change in proportion of dividing cells in S-phase (Fig 3C, F), but these results support a significant reduction in specifically the proportion of dividing Beclin1-null NPCs in S-phase of the cell cycle.

### Beclin1-null cells regulate cell cycle exit differentially

To assess what changes in cell fate underlie the preferential clustering of Beclin1-null cells into the CCE3 group, we used scVelo^31^ that revealed fate choice vectors of Beclin1-null and WT proliferating cells based on mature and unprocessed RNA dynamics. As expected, the WT mitotic cells entered and largely remained in mitosis followed by entering the CCE1 and CCE2 population (Fig 5A). In contrast, the Beclin1-null cells had little mitotic activity and more robust fate vectors towards CCE3 (Fig 5B). Top 100 cluster-specific genes driving these trajectories were also revealed using scVelo’s latent time function (Suppl. Table 1). In accordance with enhanced cell death and a reduction in mitosis, we observed a loss of regulation of the Beclin1-null mitotic clusters by the apoptosis and DNA damage suppressor, Birc5/survivin^32,33^ (Suppl. Table 1, Fig 5C). Beclin1-null cells exiting cell cycle showed preferential regulation by genes such as Nfib, a negative regulator of proliferation and positive regulator of cell cycle exit^34,35^ (Fig 5D). Beclin1-null mitotic and CCE cells showed regulation by unique transcripts such as Ttc28 that regulates midzone organization during late mitosis^36^ (Fig 5E), and MAPT, a structural protein involved in cell cycle and nuclear maintenance^37^ and related to neurodegeneration^38^ (Fig 5F). Together these data reveal that Beclin1-null mitotic cells follow a differential developmental trajectory accompanied by changes in key players that regulate mitosis and differentiation, and directly activate genes impeding proliferation.

**Figure 5.**
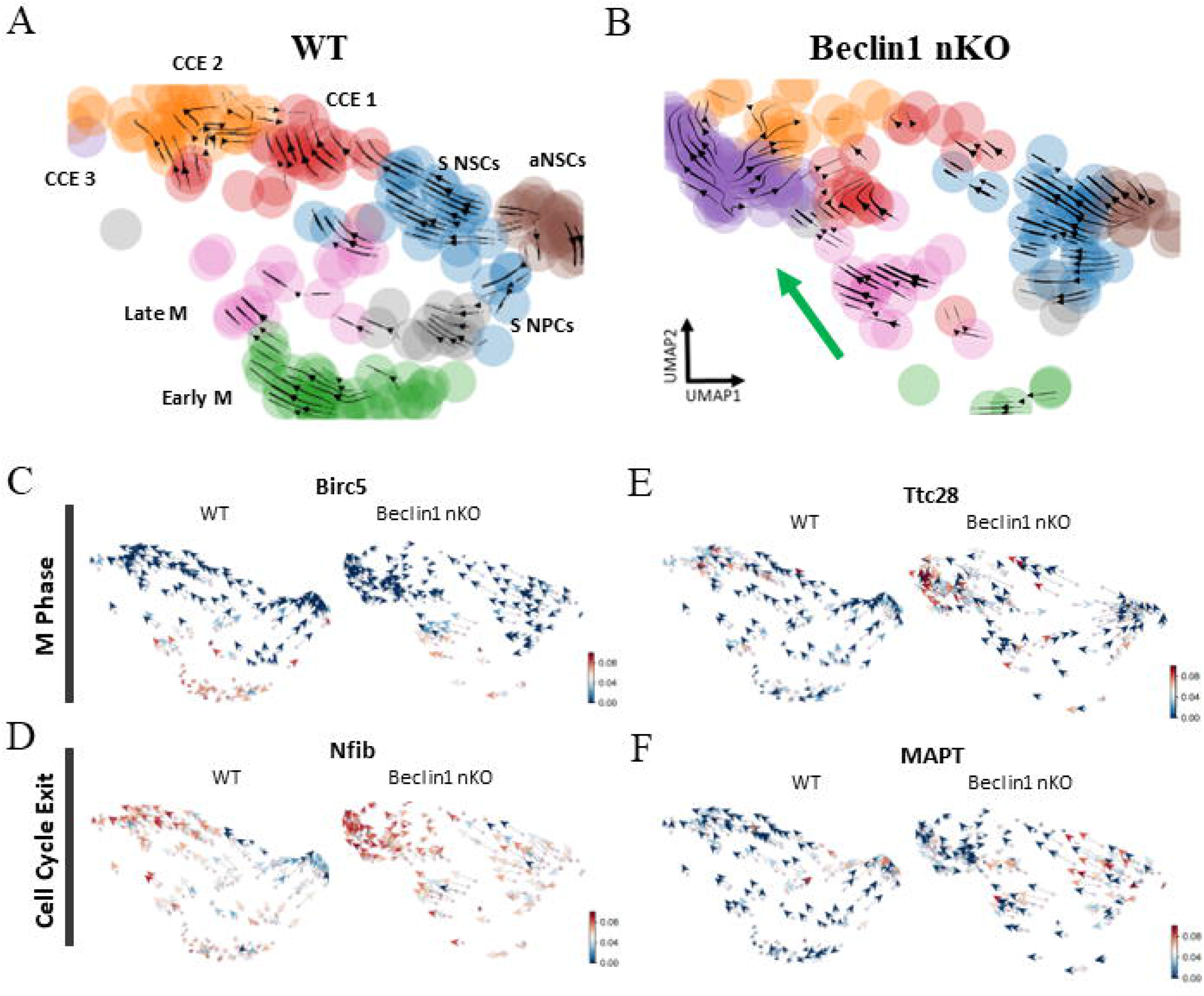
Velocity analysis of WT and Beclin1 nKO proliferating cells identifies differential transcriptional regulation during mitosis and CC exit. (A) WT and Beclin1 nKO cycling cell UMAP projections of cycling clusters with added RNA velocity vectors (black arrows) showing a high propensity of Beclin1 nKO mitotic cells toward exit into CCE3 cluster, while (B) mitotic WT cells show a differential trajectory of highly proliferative state toward CCE1/2 upon exit. M phase and CC exit cells demonstrate reduced transcriptional activity (red arrows/high activity, blue arrows/low activity) of (C) Birc5, and increased activity of (D) Nfib, (E) Ttc28, and (F) MAPT.

### Beclin1-null cells downregulate mitotic genes and upregulate cell stress genes upon cell cycle exit

There were 223 significantly differentially expressed genes (DEGs) in WT and Beclin1 nKO mice: 49 reduced genes and 174 upregulated (Suppl. Table 2). Corroborating scVelo findings of decreased mitotic transcriptional activity, Beclin1-null cells showed decreased expression in DEGs that were abundantly expressed in the mitotic WT cells. This includes genes that included parts of the mitotic machinery such as Cenpk/m/h^39,40^, Kif11^41^, Smc2^42^, that make up the centromere^39^, kinesin^43^ and condesin I and II^44,45^ complexes, respectively, which play roles in the preparation for cell division and in chromatin maintenance^46–48^ (Fig 6A; Suppl. Table 2). Furthermore, dysregulation of the kinetochore was evident in Beclin1-null cells by the reduced expression of Kif22^49^, Spc24 and Spc25^50^ (Suppl. Table 2), as well as Ran and RanBP1 that are required for proper attachment of kinetochores to microtubules^51–53^ (Suppl. Table 2). There was also decreased expression of DNA repair genes such as Rrm2^54^ and Ran GTPase^51,55^ (Fig 6A), as well as others such as Top2a ^56^, Pclaf ^57^, and Syce2 ^58^ (Suppl. Table 2). In contrast, the genes that had enhanced expression in Beclin1-null cells were cell stress, metabolism, and neurodevelopment genes, such as Tpt1 ^59^, Tmxb4x ^60^, Snapin ^61^, Nptn ^62^, Tcf4 ^63^, and Cplx2 ^64^ (Fig 6B). Additionally, Beclin1-null cells showed upregulation in cell stress-related transcripts such as reactive oxygen species genes (ROS; e.g., Romo1 ^65^, Pet100 ^66^, Ndufa3 ^67^; Suppl. Table 2), and P53 targets (e.g., Tpt1 ^68,69^, Cox2 ^70^, Sesn1 ^71^; Suppl. Table 2). These genes also had more enhanced and specific expression in the CCE3 cluster in Beclin1-null cells compared to WT cells.

**Figure 6.**
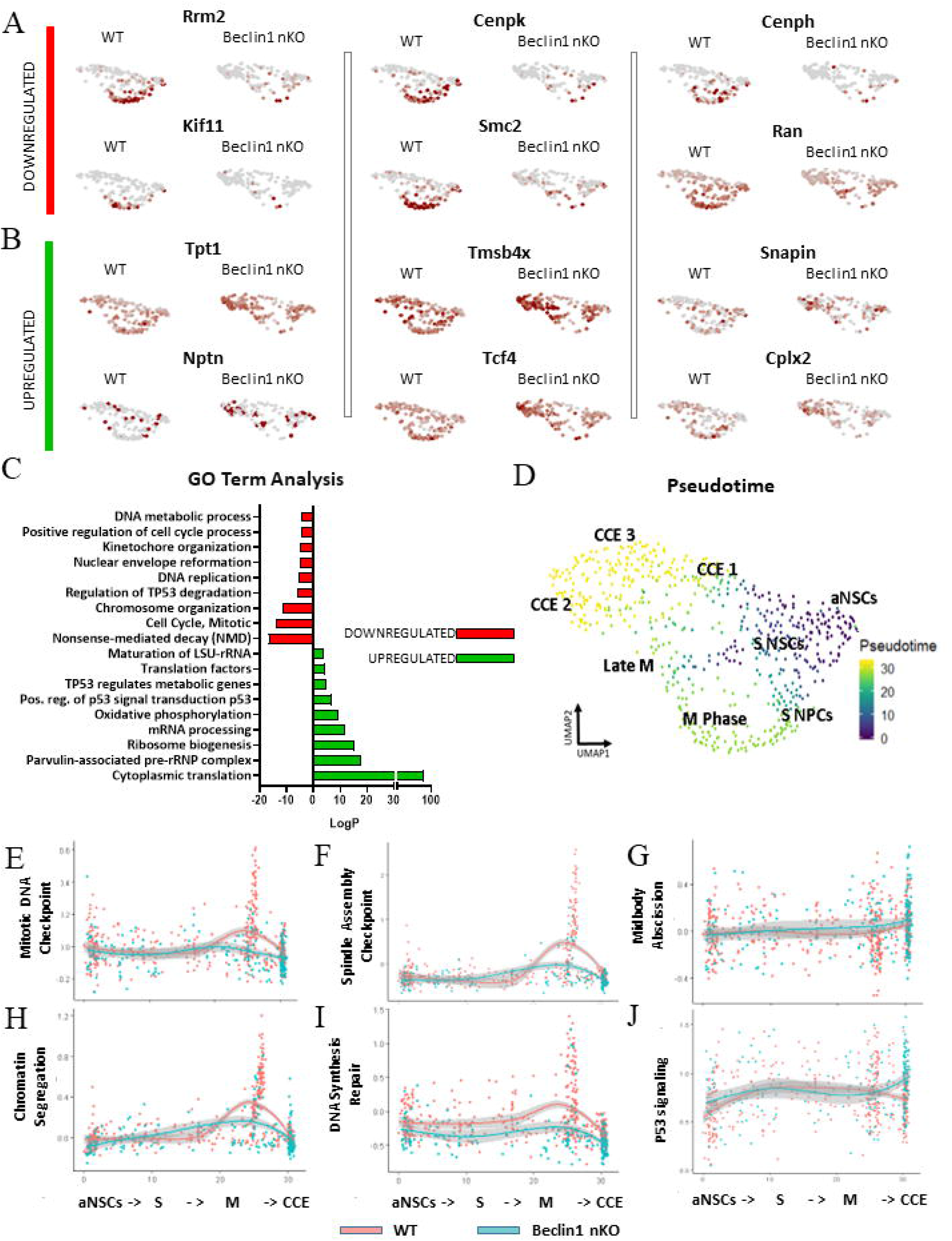
Beclin1-null NSPCs differentially downregulate mitotic maintenance genes and increase transcription of cell stress signatures. (A) WT and Beclin1 nKO cycling cell UMAP projections of relative RNA expression of genes downregulated in the Beclin1 nKO sample showing localization in mitotic cells. (B) WT and Beclin1 nKO cycling cell UMAP projections of relative RNA expression of genes upregulated in the Beclin1 nKO sample showing localization in cells exiting cell cycle. (C) Gene ontology analysis results shown as LogP values (LogP≥4) for processes upregulated (green) and downregulated (red) in Beclin1 nKO cycling cells. (D) UMAP projection of Pseudotime analysis of cycling WT and Beclin1 nKO cells showing a gradient from relative time 0 (indigo) to relative time 30 (yellow). Combined relative RNA expression of GO Terms along pseudotime showing cell pathways active in WT and Beclin1 nKO cell throughout cell cycle with reduction (E) mitotic DNA checkpoint and (F) spindle assembly checkpoint genes, with (G) no change in midbody abscission. Deficits were also observed in (H) chromatin segregation, (I) DNA synthesis repair, and (J) p53 signaling genes. Individual dots represent cells, lines represent mean calculated expression values for GO Term genes across pseudotime, filled gray line shows standard deviation.

Gene ontology (GO) term analysis of the upregulated and downregulated differentially expressed genes further revealed that cellular processes relating to cytoplasmic translation, ribonucleoprotein and ribosome generation, oxidative phosphorylation, and p53 signaling were significantly enhanced in Beclin1-null proliferating cells (Fig 6C). The cells also had a decreased expression of processes relating to nonsense-mediated decay of RNA, mitotic cell cycle, chromosome and kinetochore organization, p53 degradation, nuclear maintenance, as well as DNA replication and metabolism (Fig 6C).

In order to observe these pathways during proliferation in relative time, we constructed pseudotime trajectories of WT and Beclin1-null cells (Fig 6D) followed by quantification of pathway activity over time using GO terms. Beclin1-null cells showed a dramatic reduction in the activity of genes necessary for the G2/M transition checkpoint that ensures DNA integrity and cell size (Fig 6E). Further on, mid-mitosis cells also failed to activate genes necessary for the spindle assembly checkpoint (Fig 6F) that ensure proper DNA maintenance in preparation for anaphase and telophase. This did not affect the regulation of cytokinesis (Fig 6G). However, the chromosome segregation genes were significantly inhibited (Fig 6H) in Beclin1-null mitotic cells together with insufficient activation of DNA repair genes (Fig 6I), suggesting that cells may accumulate DNA damage overtime. Lastly, Beclin1-null cells had enhanced p53 signaling at mitotic exit (Fig 6J) which could mediate apoptosis. Together these data corroborate our findings *in/ex vivo* and *in vitro* and suggest that deficits in chromatin dynamics in Beclin1-null cells reduces the mitotic activity and increases death of the daughter cells upon cell cycle exit.

### Beclin1 nKO mice have accumulation of DNA damage and fewer adult-generated granule neurons

The faulty mitotic checkpoints and DNA maintenance that were seen in Beclin1-null cells can commonly result in DNA damage and chromatin abnormalities such as aneuploidy, genomic instability, micronuclei, and lagging chromosomes in cancer cells^72,73^. To determine if Beclin1-null cells have reduced viability due to DNA damage accumulation, we tested the general levels of fragmented DNA in an alkaline comet assay that is highly sensitive to both double- and single-strand DNA breaks^74^. Sorted Beclin1-null cells had a higher percentage of DNA in the comet tail (Fig 7A, B) suggesting increased DNA fragmentation and supporting the notion that Beclin1-null cells have increased accumulation of DNA damage.

**Figure 7.**
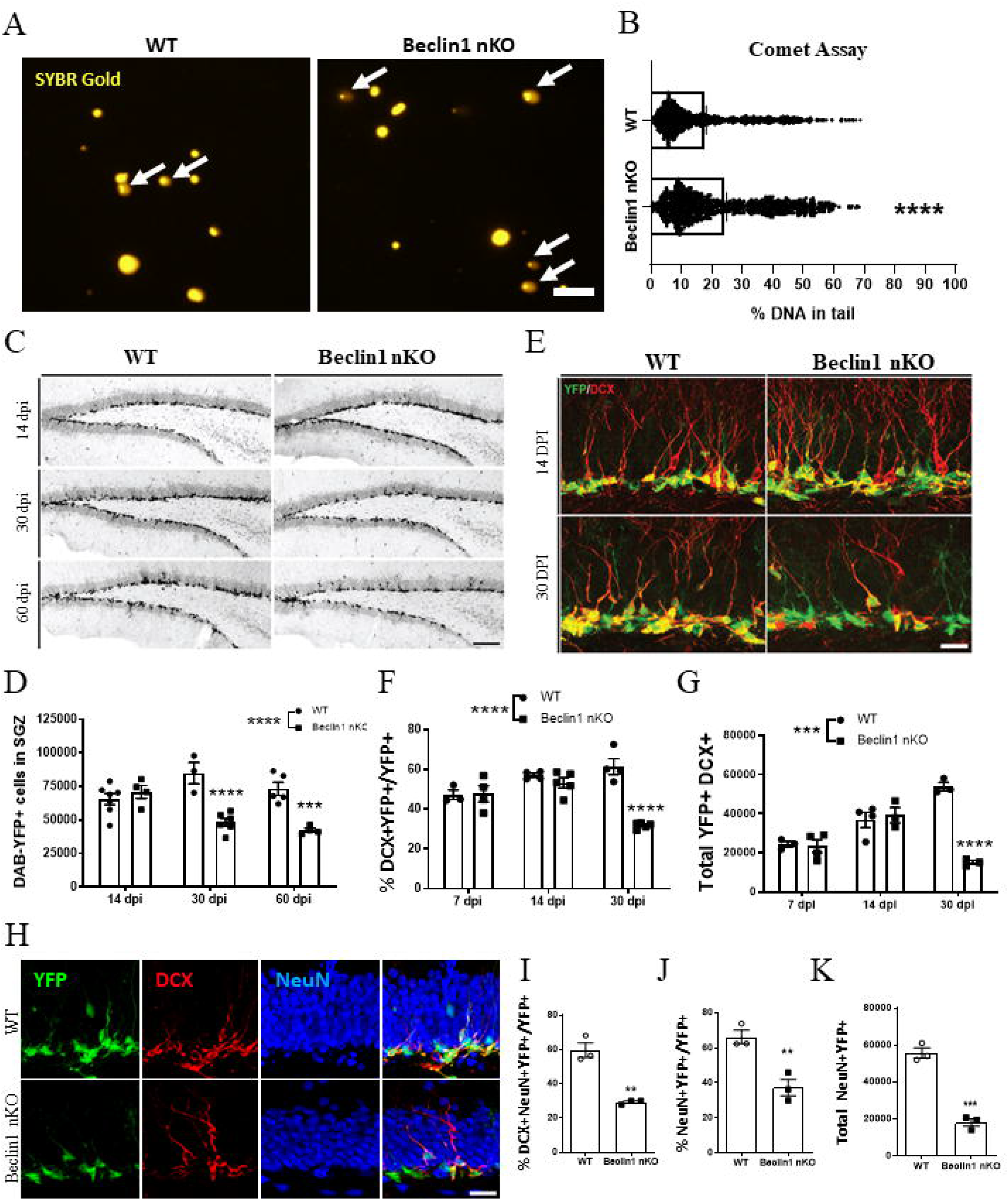
Beclin1 removal increases DNA damage, reduces hippocampal neurogenesis and survival. (A) Representative fluorescent images and quantification (B) of DNA damage visualized using SYBR-Gold (yellow) in WT and Beclin1 nKO recombined cells sorted at 14 dpi showing fragmented DNA (A, white arrows) and (B) a significant increase of DNA tail percentage in Beclin1 nKO samples. (C) Representative DAB-YFP images and (D) quantification the YFP+ cells (black) in the dentate gyrus in WT and Beclin1 nKO mice showing a reduction YFP+ cells at 30 and 60 dpi (E) Representative images of the immature neuronal marker doublecortin (DCX, red) and YFP+ (green) cells shows a reduction in (F) percentage of YFP+ that expressed DCX and (G) total number of DCX+YFP+ neurons in the Beclin1 nKO mice at 30 dpi. (H) Representative images of the YFP (green), DCX (red) and mature neuronal marker (NeuN). blue) cells shows a reduction in the percentage of (I) maturing (DCX+NeuN+YFP+) and mature neurons (NeuN+,YFP+) as well as (K) total of mature neurons in Beclin1 nKO mice. Scale bars represent 20 uM (A); 500 uM (C), 25 uM (E), 25 uM (H); All graphed data show individual animals (D-G, I-K), or cells (B), as well as the mean ± SEM; Data with 2 groups were analyzed by unpaired t-test, data with more than 2 groups were analyzed by a 2-way ANOVA; ** p≤0.01, *** p≤0.001, **** p≤0.0001.

To test if the accumulation of DNA damage results in the reduction in the generation of adult-born cells, we performed lineage-tracing of the Beclin1-null and WT NSPCs and their progeny between 14 to 60 dpi. There was a significant reduction in the number of YFP+ cells in the Beclin1 nKO mice compared to WT mice that started at 30 dpi (Fig 7C, D). Since the reduction in number of Beclin1-null cells started 30 days after removal of Beclin1, we tested if Beclin1 was reducing the survival of adult-generated neurons, in addition to reducing proliferation. Through using a retroviral-based approach we have previously published to track the survival of the late-stage dividing NPCs^14^, we found no difference in survival of the NPCs at 14, 30, and 60 days after removal of Beclin1, with small but significant overall reduction in survival ratio in the floxed Beclin1 compared to WT mice (Fig S3A, B). Additionally at 60 dpi, there were no significant differences in spine density between the Beclin1-null and WT cells (Fig S3C, D). Given that the overall reduction in survival of floxed Beclin1 late-stage cells was minor, in comparison to the robust reduction proliferation in the Beclin1 nKO, Beclin1 more likely reduces the number of immature neurons due the reduction in proliferation, which then resulted in decreased survival of daughter cells.

To determine if the reduction in the number of Beclin1-null cells in the Beclin1 nKO mice reduced the number of adult-generated neurons in the dentate gyrus, the population of cells expressing the immature neuronal marker DCX and mature neuron marker NeuN was quantified. The Beclin1 nKO mice compared to WT mice had a significant overall reduction in number of DCX+ cells starting at 30 dpi (Fig S4A, B). There was also a significant reduction in the proportion of YFP+ cells that expressed DCX in the Beclin1 nKO mice compared to controls (Fig 7E-G). Further examination revealed a significant reduction in the proportion of late immature neurons (DCX+) that expressed the mature neuronal marker NeuN (Fig 7H, I), as well as the overall proportion and number of Beclin1-null cell that expressed NeuN (Fig 7H, J, K). Overall, these results support that a lack of Beclin1 induces changes in chromosomal dynamics during mitosis that cause DNA damage and results in the reduction in the generation of adult-born neurons.

## DISCUSSION

The results of this study demonstrate that Beclin1 is required cell-autonomously for the proliferation and mitosis of adult NSPCs to sustain adult hippocampal neurogenesis. Our data led to the novel discovery that mitosis in the adult NSPCs is regulated by Beclin1. Mitotic Belcin1-null cells had a dysregulation of genes that are necessary for chromosome maintenance during prophase and metaphase, highlighting that chromatin dynamics during cell cycle in NSPCs is regulated by Beclin1. In addition, dividing Beclin1-null cells had enhanced transcription of genes related to metabolic stress and reduced DNA maintenance that is accompanied by DNA damage that ultimately leads to cell death and a reduction in the number of adult-born granular neurons in the hippocampus.

Our findings support that Beclin1 is required in adult neurogenesis, as first identified in the SVZ of heterozygous adult Beclin1 knockout mice^12^. Similarly our findings align with the improvement in age-related reductions of NSCs in *Becn1^F^*^121^*^A/F^*^121^*^A^* knockin mice^75^. Our findings also extend these observations by discovering that Beclin1 regulates adult neurogenesis by regulating mitosis. Our *in vitro* live imaging results reveal Beclin1 is required to sustain consecutive mitotic divisions and the survival of NSPCs. This finding parallels our *in vivo* results illustrating deficits in proliferation and mitosis occurring two weeks after Beclin1 was removed, and two weeks before the reduction in number of immature neurons and adult-generated neurons.

Our findings further show that the reduction in autolysosomes accompanies the deficits in adult neurogenesis. These results align with the essential role for Beclin1 in initiation of the autophagosome^1,82^, and the reduction in autophagy flux shown following removal of Beclin1^11,13,84^. These data suggest that the actions of Beclin1 in mitosis in NSPCs rely on Beclin1’s autophagy-dependent mechanisms yet do not exclude the possibility that the effects of Beclin1 could occur independent of autophagic mechanisms. Indeed, studies examining different autophagy proteins have highlighted that not all autophagy proteins regulate neurogenesis in development and the postnatal brain^85^. Notably, the embryonic removal of Atg16L^86^ or embryonic and adult removal of Atg7^86,87^, produced no deficits in proliferation or the generation of adult born granule neurons. In contrast, our previous work suggested that autophagy–dependent mechanisms regulate neurogenesis due to the delayed survival of adult-generated neurons following retroviral-mediated deletion of Atg5 from NPCs^14^. Similarly, a significant reduction in survival of adult-generated neurons, as well as depletion of the stem cell pool occurred following NSPC-specific removal of FoxO3^88^, which is involved in the formation of the phagophore. A combination of autophagy-dependent and -independent mechanisms have been suggested in studies examining the requirement of FIP200 (focal adhesion kinase family interacting protein of 200 kD), which recruits the autophagic machinery. Specifically, mice with conditional embryonic deletion of FIP200 at 4 weeks postnatal^89^ had autophagy-dependent deficits in proliferation^90^ as well as autophagy-independent deficits in differentiation^91^. Together these studies highlight the diversity in effects of autophagy related proteins in adult neurogenesis associated with the growing appreciation for autophagy-independent functions of these proteins.

Mechanisms that regulate the cell cycle in NSPCs are beginning to be elucidated and our findings reveal Beclin1 as a specific regulator of mitosis. These findings support the work of others that reveals *in vitro* deletion of Beclin1 induces chromatin abnormalities during the cell cycle via the deregulation of the mitotic machinery^10,76,77^. The ability of our analysis to separate the proliferating NSPCs into different stages of the cell cycle further revealed Beclin1 is required in regulating differential expression of mitotic spindle, centromere, and kinetochore transcripts. For example, Beclin1 loss dysregulated expression of Cenp family genes, chromosomal condensins, and genes sustaining the kinetochore and microtubule attachment.

A failure to undergo mitosis can induce DNA damage^73^ and our findings show that Beclin1-null proliferating NSPCs have fragmented DNA. These findings align with the connections between Beclin1 and DNA repair/damage in tumorigenesis^78,79^, as well as the direct interactions of Beclin1 with DNA repair and anti-apoptotic genes^1,9,80,81^. We show that Beclin1 is required in adult NCPSs to regulate transcriptional activity of genes involved in metabolic stress, as well as DNA repair and pro-survival genes. Specifically, we showed upregulation of genes underlying reactive oxygen species response, P53 target genes, and a reduction in DNA synthesis and repair genes. For example, neurodevelopmental gene and binding partner of Beclin1 Birc5/Survivin^32,82,83^ had reduced transcriptional activity in Beclin1-null cells, and its deficiency is associated with apoptosis via mitotic deficit and DNA damage in different cell lines^33^. These findings demonstrate that under normal physiological conditions Beclin1 sustains mitosis and DNA maintenance by regulating the expression of DNA repair and metabolic stress genes.

The findings from this study demonstrate the requirement of Beclin1 in the maintenance of mitosis of adult neurogenic cells to produce granular neurons in the dentate. Beclin1 removal resulted in a reduction of mitotic cells mediated by transcriptional dysregulation of genes important for genomic maintenance during metaphase and anaphase. These effects were accompanied by increased signatures of cellular stress upon cell cycle exit, and accumulation of DNA damage. Overall, this study links the reduction of Beclin1 in dividing NSPCs with mitotic chromosome maintenance and DNA damage and thus identifies an intrinsic regulator of mitosis within the adult NSPCs.

## Acknowledgements

The authors would like to acknowledge the assistance and use of the: Flow Cytometry & Virometry (FCV) Core Facility at the University of Ottawa; Flow Cytometry & Cell Sorting facility (OHRI; RRID:SCR_023349); StemCore Laboratories Genomics Core Facility at the Ontario Health Research Institute (RRID: SCR_012601); and Cell Biology and Image Acquisition Core (RRID: SCR_021845) funded by the University of Ottawa, Ottawa, Natural Sciences and engineering Research Council of Canada, and the Canada Foundation for Innovation. We also acknowledge and thank Dr. Ryoichiro Kageyama and Dr. Zhenyu Yue for sharing their transgenic mouse lines.

## STAR Methods

### KEY RESOURCES TABLE

| REAGENT or RESOURCE | SOURCE | IDENTIFIER |
| --- | --- | --- |
| Antibodies |  |  |
| Chicken Anti-GFP (1:5000) | Aves | GFP-1020 |
| Living Colors Rabbit Anti-DsRed Polyclonal Antibody (1:500) | Clontech | 632496 |
| Goat Anti-Doublecortin (C-18) (1:500) | Santa Cruz | SC8066 |
| Rabbit Anti-Ki67 Monoclonal Antibody (1:200) | Cell Marque | 275R-14 |
| Mouse anti-BM28 (Mcm2; 1:200) | BD Transduction Laboratories | 610701 |
| Mouse Anti-NeuN Clone A60 (1:500) | Millipore | MAB377 |
| Rabbit anti-Aldolase C (1:1000) | Novus Biologicals | NBP1-90954 |
| Goat anti-Sox2 (1:1000) | Neuromics | GT15098 |
| Rabbit anti-Beclin1 (1:250) | Santa Cruz | SC11427 |
| Biotin-SP-AffiniPure Donkey Anti-Chicken IgY (IgG) (1:200) | Jackson Laboratories | 703-065-155 |
| Biotin-SP-AffiniPure Donkey Anti-Rabbit IgG (1:200) | Jackson Laboratories | 711-065-152 |
| Biotin-SP-AffiniPure Donkey Anti-Goat IgG (1:200) | Jackson Laboratories | 705-065-147 |
| Alexa Fluor 488 AffiniPure F(ab')<sub>2</sub> Fragment Donkey Anti-Chicken IgY (IgG) (H+L) (1:500) | Jackson Laboratories | 703-546-155 |
| Alexa Fluor 594 AffiniPure F(ab')<sub>2</sub> Fragment Donkey Anti-Goat IgG (H+L) (1:500) | Jackson Laboratories | 705-586-147 |
| DyLight649 AffiniPure F(ab')<sub>2</sub> Fragment Donkey Anti-Rabbit IgG (H+L) (1:500) | Jackson Laboratories | 711-495-152 |
| Alexa Fluor 647 AffiniPure F(ab')<sub>2</sub> Fragment Donkey Anti-Goat IgG (H+L) (1:500) | Jackson Laboratories | 705-606-147 |
| Alexa Fluor 594 AffiniPure F(ab')<sub>2</sub> Fragment Donkey Anti-Mouse IgG (H+L) (1:500) | Jackson Laboratories | 715-586-150 |
| Alexa Fluor 594 AffiniPure F(ab')<sub>2</sub> Fragment Donkey Anti-Rabbit IgG (H+L) (1:500) | Jackson Laboratories | 711-585-152 |
| Cy3 AffiniPure F(ab')<sub>2</sub> Fragment Donkey Anti-Mouse IgG (H+L) (1:500) | Jackson Laboratories | 715-165150 |
| Cy5 AffiniPure F(ab')<sub>2</sub> Fragment Donkey Anti-Rabbit IgG (H+L) (1:500) | Jackson Laboratories | 711-175-152 |
| Chemicals, peptides, and recombinant proteins |  |  |
| TrypLE Express | Gibco | 12604021 |
| Fixation Buffer | Biolegend | 420801 |
| Intracellular Staining Permeabilization Wash Buffer (10X) | Biolegend | 421002 |
| SYBR™ Gold Nucleic Acid Gel Stain | Invitrogen | S11494 |
| Sulfo-Cyanine3 Azide | Lumiprobe | A1330 |
| Sulfo-Cyanine5 Azide | Lumiprobe | A3330 |
| 5-Ethynyl-20-deoxyuridine | Santa Cruz | sc-284628 |
| Tamoxifen | Sigma | T5648-5G |
| Sunflower seed oil from *Helianthus annuus* | Sigma | S5007-250ML |
| DMEM-F12 no phenol | Gibco | 11039-021 |
| DMEM-F12 with phenol | Gibco | 11330032 |
| Penicillin-streptomycin | Multicell | 450-201-EL |
| Ply-d-lysine hydrobromide | Sigma | P0899-10MG |
| Percoll 100% | GE Healthcare Life Sciences | 17-0891-02 |
| Papain | Worthington Biochemical Corp. | LS003126 |
| Fibroblast growth factor (FGF) | Sigma Aldrich | F0291-25UG |
| Epidermal growth factor (EGF) | Promega | 39172901 |
| B-27 Serum-Free Supplement (50X) (10ml) | Gibco | 17504044 |
| 7-Amino-Actinomycin D (7AAD) | Sigma | A9400-1MG |
| Critical commercial assays |  |  |
| Metal Enhanced DAB substrate kit | Thermo Scientific | 34065 |
| ABC elite kit | Vector Laboratories | PK-6100 |
| Deposited data |  |  |
| sc-RNAseq data | This study | GSE247493 |
| Code to analyze sc-RNAseq data | This study |  |
| Software and algorithms |  |  |
| R software | R Core Team (2021) |  |
| Python | Python Software Foundation |  |
| Seurat | Stuart et al. (2019)<sup>92</sup> |  |
| Monocle | Trapnell et al. (2014)<sup>93</sup> |  |
| scVelo | Bergen et al. (2020)<sup>94</sup> |  |
| FlowJo | Becton, Dickinson and Company |  |
| Zen Blue/Black | GraphPad Software |  |
| ImageJ/Fiji | ImageJ |  |
| GraphPad Prism | Zeiss group |  |
| Columbus Image Data Storage and Analysis System | Perkin Elmer |  |
| BioRender (graphical abstract) | BioRender.com |  |

### CONTACT FOR REAGENT AND RESOURCE SHARING

Further information and requests for resources and reagents should be directed to and will be fulfilled by the Lead Contact, Dr. Diane Lagace.

### EXPERIMENTAL MODEL AND METHOD DETAILS

#### Animals

Animal procedures were performed with approval from the University of Ottawa Animal Care Committee and adhered to the Guidelines of the Canadian Council on Animal Care. Beclin1 nestin-inducible knockout transgenic mice (Beclin1 nKO) were created by breeding using: inducible Nestin-CreERT2 mice^95^ (line 4.1 obtained from Dr. Ryoichiro Kageyama, Kyoto University, Japan); floxed Beclin1 (fBeclin1) mice (obtained from Dr. Zhenyu Yue, Icahn School of Medicine at Mount Sinai, USA)^13^; and reporter R26R-enhanced Yellow Fluorescent Protein (YFP) mice^96^ (Jackson Laboratory). These mice allow for the conditional removal of Beclin1 and expression of YFP from nestin-expressing stem and progenitor cells, as well as all of their progeny following injection of tamoxifen (TAM). Experimental Beclin1 nKO mice were NestinCreERT2^het^;RosaYFP^het;^fBeclin1^homo^ and wild-type (WT) control mice were NestinCreERT2^het^;RosaYFP^het;^fBeclin1^WT^. Age-, sex- and littermate-matched Beclin1 nKO and WT mice were randomly used as experimental mice based on their genotype. All strains were obtained and maintained on a C57bl/6J background. Animals were group housed in standard laboratory cages and kept on a 12-hour night/day cycle with ad libitum access to food and water.

#### Genotyping

Animals were genotyped at 3 weeks of age through DNA samples obtained from ear clippings (∼1 mm^2^). DNA was extracted using the HotSHOT methodology^97^. Briefly, ear clippings were incubated an Alkaline Lysis Buffer (25 nM NaOH and 0.2 mM Na2EDTA) at 95°C for 30 minutes prior to addition of the Neutralization Solution (40 mM Tris-HCl). Polymerase Chain Reaction (PCR) was completed using primers (Fig S5A) according to previously published protocols for fBeclin1^13^, Nestin-CreERT2^95^, and R26R-eYFP^96^. The resulting PCR products were resolved by size on a 1% agarose gel using electrophoresis. Size of the PCR products was visualized with ethidium bromide staining under ultraviolet light and estimated by comparison with a 100 base pair (bp) DNA ladder (DM001-R500M; Frogga Inc).

#### Tamoxifen Administration

Tamoxifen (TAM, T5648-5G; Sigma) was administered via intraperitoneal (IP) injection at a dosage of 160 mg/kg/day for 5 days (dissolved in 10% EtOH and 90% sunflower oil) to 4-8 week-old Beclin1 nKO and control mice, similar to previously published work^98^. For all experimental time points (7, 14, 30, and 60 days post injection of TAM) a minimum of 3 animals per genotype were analyzed.

#### EdU Administration

5-Ethynyl-2′-Deoxyuridine (EdU; sc-284628; Santa Cruz) was given to the mice through IP injection at a dosage of 50 mg/kg. Mice were injected with EdU four times two hours apart and perfused 8 or 24 hours after the first injection in order to obtain slices for immunohistochemistry (IHC). Similarly, for flow cytometry and cell cycle analyses, mice were injected with EdU four times two hours apart and tissue was harvested 24 hours after the first injection.

#### Preparation and Stereotaxic Surgical Injection of Retroviruses

To visualize the autolysosome within the brain, the mCherry-EGFP-LC3 retrovirus was created as we have published^14^ using the mCherry-EGFP-LC3B plasmid (Addgene, Cambridge, MA, USA, 22418). The retrovirus (1.5µl volume) was injected bilaterally into the dentate gyrus of Beclin1 nKO or WT mice (7–9 weeks old) during stereotaxic surgery 3 days after TAM injections.

To visualize the survival of dividing progenitor cells the *GFP-Cre* and *RFP* retroviruses were created using the retroviral vectors *CAG-GFP-Cre* and *CAG-RFP* and corresponding packing envelopes, that were generously provided from Dr. Fred Gage (Salk Institute of Biological Science). Either *GFP-Cre* and *RFP* retroviruses in a 1:1 ratio mixture (volume 1.5 μl) or *GFP-Cre* (volume 1 μl) retrovirus were injected bilaterally into the dentate gyrus of 7-9 week old floxed Beclin1 (fBeclin1) (homo, or WT control) ^13^ mice using stereotaxic surgery. The viruses were created using a previously published protocol^99^ with minor modifications. Briefly, 293T cells were plated (8×10^6^ cells/150 mm) and co-transfected using polyethylenimine (PEI; 23966; Polyscience) with either the *CAG-GFP-Cre* or *CAG-RFP* retroviral plasmid combined with the *CMV-Gag-Pol* packing plasmid and *CMV-VSV-G* envelope plasmid in a 3:2:1 ratio, respectively. At 48 and 72 hours post-transfection the supernatant containing the virus was collected and concentrated by two rounds of ultracentrifugation (20,000 RPM for 2 hours at 4°C) with 20% sucrose cushion, dissolved in phosphate buffered saline (PBS). Virus titre was determined by live tittering through infection of 293T cells plated in a 24-well plate (1.25 x 10^5^ cells) with 100 μl of diluted (10^4^ dilution) virus. Fluorescence-positive cells were quantified 48 hours post-infection and the number of infectious units (IU) per ml was calculated as the mean of the product of the number of infected cells per viewing field, the well area (243.22 mm2), and the dilution factor (104). Virus titre was approximately 6.7 x 10^8^ IU/ml for the *GFP-Cre* virus and 1.7 x 10^9^ IU/ml for the *RFP* virus.

During stereotaxic injection, the mice were anesthetized throughout surgery with 2% isoflurane. Viral injections were performed by microinjection using a 33 gauge (0.21 mm diameter) needle (7803-05; Hamilton), into the dentate gyrus using coordinates of -1.7 mm rostrocaudal and ±1.2 mm mediolateral from bregma, and -2.4 mm dorsoventral form the skull surface. The virus was injected using a Nanomite Pump 11 Elite (704507; Harvard Apparatus) at a rate of 0.2 μl/min and the needle was removed 5 minutes after the injection was complete in order to prevent backflow. Post-operation recovery from anesthesia occurred in a 37°C incubator until mice were awake and responsive. Buprenorphine was given to the mice as an analgesic (0.05 mg/kg, subcutaneous injection) one hour before surgery, as well as 6 and 12 hours after viral injection. The mice were sacrificed and perfused at 17 days after viral injection of the mCherry-EGFP-LC3, and after 14, 30, or 60 days after dual *GFP-Cre:RFP* retrovirus delivery.

#### Western Blot

Tissue from Beclin1 nKO or WT mice were dissected 35 days after Tam treatment. In order to harvest a higher number of cells for proteins lysates the tissue was dissected from adult subventricular zone which contains more YFP+ cells compared to the SGZ^100,101^, cells were then incubated in a digestion media containing DMEM/F12 (11039-021; Invitrogen), 1.2 mM EDTA (E5134-1KG; Sigma), and 20 U/ml papain (LS003126; Worthington Biochemical) at 37°C for 30 min. Cells were triturated followed by centrifugation to obtain a cell pellet that was suspended in media containing DMEM/F12 and 10% Fetal Bovine Serum (FBS; SH3039603; HyClone) to inactivate papain. YFP+ and YFP-cells negative for 7-AAD were then sorted using MoFlo Astriosis (Beckman Coulter). Sorted YFP+ and YFP-cells were lysed in 8 mM urea with 10% sodium dodecyl sulfate (SDS). The lysed samples were mixed with an equal volume of laemmli loading buffer with 10% b-mercaptoethanol, boiled at 95°C, vortexed, and loaded onto a 12% acrylamide gel. The gel was immersed in 1X tris/glycine/SDS (TGS) running buffer and run at 110V for 1.5 hours for optimal band separation. Bands were transferred to a nitrocellulose membrane via a wet transfer in cold 1X Tris/Glycine transfer buffer containing 20% methanol for 1 hour at 110V. The nitrocellulose member was cut into two for detection of Beclin1 (60 kDa) and HistoneH3 (18 kDa). The blots were incubated for 1 hour at room temperature (RT) in a blocking solution containing 5% non-fat dried milk in 1X TBS-T (0.1% Tween-20 in 1X TBS) followed by incubation in blocking solution containing either the primary antibody for Beclin1 (1:1000; sc11427; Santa Cruz) or HistoneH3 (1:1000, ab1791, Abcam) overnight at 4°C. The following day at RT the blots were washed with TBS-T and incubated for 1 hour in blocking solution containing corresponding horseradish peroxidase conjugated secondary antibodies (1:5000). After secondary incubation, the blots were washed in TBS-T incubated in ECL Pierce for 5 minutes to allow chemiluminescence detection. The blot was imaged using a Fuji LAS-4000mini chemiluminescence imager (Thermo Fisher Scientific) and densitometry was performed using Fiji image processing software (ImageJ) to determine relative amounts of protein.

#### Perfusions and Tissue Collection

Mice were anesthetized with euthanyl (90 mg/kg) and transcardially perfused with cold 1X phosphate buffer solution (PBS, pH 7.4) for 6 minutes and subsequently cold 4% paraformaldehyde (PFA) in 1X PBS (pH 7.4) for 15 minutes at a rate of 7 ml/minute. Brains were removed and postfixed in 4% PFA for 1 hour and then transferred to 30% sucrose in 1X PBS for cryoprotection. Brains were coronally sectioned into 25 or 30 μm slices with a freezing microtome (Leica SM 2000R, Leica Microsystems) and stored in PBS with 0.1% sodium azide.

#### Antibodies and Immunohistochemistry (IHC)

All primary and secondary antibodies as well as their concentrations used for immunohistochemistry (IHC) are listed in STAR methods table. Notably, a Green Fluorescent Protein (GFP) primary chicken antibody was used to detect both YFP immunoreactive (YFP+) cells in the Beclin1 nKO mice, and GFP-Cre (GFP+) cells in the virally injected fBeclin1 mice.

Slide-mounted IHC was used to detect the total number of YFP+ cells, DCX+ cells, and Ki67+ cells within the SGZ using previously published protocols ^98,102^. Briefly, every ninth section through the mouse hippocampus was mounted onto charged slides and allowed to dry overnight. Slides were then pre-treated to enhance antigen retrieval with 0.1M citric acid (pH 6.0) at approximately 95°C for 15 minutes, rinsed in 1X tris-buffer saline (TBS), incubated at RT in 0.1% trypsin for 10 minutes, rinsed in 1X TBS and then incubated with 2N hydrochloric acid (HCl) at RT for 30 minutes. To prevent non-specific binding, the slides were incubated in 3% Normal Donkey Serum (NDS; 017-000-121; Jackson Immuno Research Laboratories Inc) in 0.3% Triton X-100 in 1X TBS for 60 minutes. Sections were then incubated overnight in the primary antibody in 3% NDS in 0.3% Tween20 and 1X TBS. The following day, slides were incubated at RT in: 1) biotinylated attached secondary antibodies in 1.5% NDS in 1X TBS for 60 minutes; 2) 0.3% H2O2 in 1X TBS for 30 minutes to quench endogenous peroxidases; 3) Avidin-Biotin Complex Solution (ABC; PK-6100; Vector Laboratories) for 90 minutes; 4) metal enhanced 3,3’-Diaminobenzidine (1:10, DAB; 34065; Thermo Scientific,) for 10-30 minutes; and 5) fast red nuclear stain (H3403; Vector) to provide a nuclear counterstain. Between all steps, with the exception of after blocking with NDS, the slides were rinsed 3x with 1X TBS. Following staining, slides were dehydrated by consecutively immersing slides in 95% and 100% ethanol for 20 seconds, followed by CitriSolv clearing agent (22-143-975; Fisher) for 20 seconds, 1 minutes, and 5 minutes. Slides were cover-slipped with DPX mounting medium (mixture of Distyrene, Plasticizer, Xylene; 44581; Sigma).

All IHC for the co-labelling of more than one marker was completed using free-floating fluorescent IHC similarly to previously published methods by our laboratory ^98,102^. Briefly, sections were washed with 1X PBS three times, and were incubated in a carrier solution (1X PBS, 0.1% TritonX-100, 0.1% Tween20) on a shaker overnight with primary antibody at 4°C, with the exception for staining for Beclin1 protein which used an incubation of 48 hrs. The following day, the sections were incubated at RT in fluorophore-conjugated secondary antibody for 1 hour in carrier solution, washed in 1X PBS. Section being processed for EdU staining were additionally incubated after the secondary antibody in an EdU-staining cocktail containing 1M Tris (pH = 8.5), 200 mM CuSO4*H2O, 4mM of sulfo-Cyanine azide (A3330, A1330; Lumiprobe), and 1M of sodium ascorbate in water for 30 minutes. All sections were counterstained with 4’,6-diamidino-2-phenylindole (1:10000, DAPI; 11836170001; Roche) and the sections were slide mounted and cover-slipped with Immumount mounting media (2860060; Fisher Scientific).

#### Microscopy and Cellular Quantification

Counts for the number DAB+ cells or immunoreactive fluorescent cells in the SGZ were performed at 40x magnification using an Olympus BX51 fluorescent microscope. Manual counting was completed exhaustively through the SGZ by a blinded experimenter in every 9^th^ section of the hippocampus as previously published^98,102^. Final counts were estimated for the whole SGZ by taking the manual count multiplied by 9, or expressed as a ration of dual-labelled GFP+RFP+ cells over total RFP+ as previously published^99^. Quantification was further verified by an additional blinded experimenter that confirmed less than 10% variation in a minimum of 2 independent animals.

For quantification of the mCherry+ autolysosomes, the YFP+ Beclin1-null or WT cells in the SGZ were imaged with a 63x oil immersion lens with a Zeiss LSM800 AxioObserverZ1 mot Confocal Microscope at emission wavelengths of 517 and 561nm. Autolysosomes were blindly and manually quantified in every YFP+ cell within the section analyzed. Autolysosomes were counted in the cell body and cell processes using ZEN 2012 Blue (Zeiss) image processing software. In order to be counted autolysosomes had to be mCherry+ (red), be circular in shape, and be larger than any observed specs in the background. Quantification was verified by an additional blinded experimenter that confirmed less than 10% variation in 2 independent counts.

For quantification of single- and co-labeled florescent immunoreactive cells was done through imaging the DG at 40x (oil immersion) from bregma matched (positions around -2.06 to -2.30) using either coronal half-brain or full-brain sections. Images were obtained using either a 1) Zeiss LSM 510-META confocal microscope at emission wavelengths of 405, 488, 543, and 633 using either 20x, 0.8 NA, Air, Plan-Apo (DIC II) or 40x, 1.3 NA, Oil, EC Plan-Neofluar (DIC III) objectives; or 2) a Zeiss LSM800 AxioObserverZ1 mot confocal microscope (Zeiss) at emission wavelengths of 405, 488, 561, and 640nm using either 20x, 0.8 NA, Air, Plan-Apo or 40x, 1.3 NA, Oil, Plan-Apo objectives. ZEN acquisition software (Zeiss) was used for 1-2 μm optical sectioning in the Z-plane. Both single- and co-labeled cells were quantified manually from images visualized through Fiji image processing software (ImageJ) and ZEN Blue (Zeiss) image processing software. The total population of YFP+ cells that co-labeled with another marker was calculated as the product of the absolute YFP counts and the proportion co-labeled per animal.

For quantification of the Beclin1-positive puncta at 14 dpi, the YFP+ Beclin1-null or WT cells in the SGZ were imaged using the Zeiss LSM800 AxioObserverZ1 mot Confocal Microscope at emission wavelengths of 405, 488, and 640nm was used with 63x, 1.4 NA, Oil, Plan-Apo objective and optical sectioning of 0.5 μm in the Z-plane and 16x binning. Images were processed in ZEN Blue (Zeiss) image processing software for single-pixel filtering, and median correction.

For quantification of the spine density in the GFP and RFP co-labeled virally infected cells, the sections were imaged at 63x (oil immersion) with a Quorum Spinning-disk confocal microscope at emission wavelengths of 406, 490, and 561. MetaMorph automation and image acquisition software (Molecular Devices) was used to create a high resolution three-dimensional representation of spines throughout the visible dendritic arbor using 0.5 μm Z-plane optical sectioning in combination with a tile-scan module. Images were subsequently stitched and flattened in MetaMorph and exported to NeuroStudio (CNIC, Ichan School of Medicine at Mount Sinai) to measure neurite length. Spines were manually quantified from a single neurite that spanned the hippocampal molecular layer (top of the granule cell layer to the hippocampal fissure) per cell in Fiji image processing software (ImageJ). Spine density (spines/10 μm) was calculated as the quotient of the number of spines over neurite length multiplied by 10 (methods adapted from Zhao et al.^103^)

#### Neurosphere Culture

For all *in vitro* and *ex vivo* analyses of the DG previously published protocols^104,105^ were used to dissect the SGZ from mice 14 days after TAM injections. For neurosphere assay^106,107^, the mice were anesthetized with euthanyl (90 mg/kg), decapitated, and their brains were removed and placed in slushy sterile filtered Artificial Cerebrospinal Fluid (aCSF, pH =7.4), consisting of (in mM): 124 NaCl, 5 KCl, 1.3 MgCl2·6H2O, 2 CaCl2·2H2O, 26 NaHCO3, and 1X penicillin-streptomycin (10,000 U/mL; 450-201-EL; Multicell). The dentate gyrus of both hemispheres was dissected out of the brain and placed in aCSF. The tissue was gently broken up with sterile scissors. Tissue was incubated on a thermomixer (30 minutes, 37°C) in 500 uL/tube of digestion media, containing DMEM/F12 (11039-021; Invitrogen),

1.2 mM EDTA (E5134-1KG; Sigma), and 20 U/ml papain (LS003126; Worthington Biochemical) at 37°C for 30 min. Cells were triturated followed by centrifugation to obtain a cell pellet that was suspended and washed in DMEM/F12 media. Suspension was then transferred in Percoll media, consisting of 19.8% Percoll (17-0891-02; GE Healthcare Life Sciences), 2.2% 10xPBS (311-012-CL; Multicell) and spun down (500 x g, 13 minutes, RT). Cells were again washed with DMEM/F12 and quantified using Countess II (Invitrogen) cell counting charged slides by reconstituting cell suspension with trypan blue at a ratio of 1:1. Cells were then seeded at 20k cells/1ml with growth media containing DMEM/F12, 1X B-27 supplement (17504044; Gibco), 1X Penicillin-Streptomycin, Heparin (H3149-25KU; Sigma), 200 ng/μl of Epidermal Growth Factor (EGF; 39172901; Promega), and 100 ng/μl of Fibroblast Growth Factor (FGF; F0291-25UG; Sigma Aldrich). Neurospheres were quantified live under an inverted Zeiss AxioObserver D1 microscope (Zeiss) using 20x, 0.80 NA, Air, Plan Apochromat (DIC II) objective and AxioCam MRm CCD as a total number of spheres per well and size was calculated using AxioVision 4.8 (Zeiss).

#### Live *in vitro* imaging

The cells used to perform using live in vitro image were removed, digested, and processed from the Beclin1 nKO and WT mice as described using the same protocol as listed above for the neurosphere assay up to the stage of cell quantification using the Countess II. After counting, for *in vitro* imaging the cells were seeded at a concentration of 40k/ml into 96-well PhenoPlates (6055302, PerkinElmer Health Sciences Canada Inc) pre-coated with Poly-D-lysine hydrobromide (P0899-10MG; Sigma), and cultured in an incubator for 5 days. On day 5 of culture the plate was sealed with Axygen™ Microplate Sealing film (14-222-346, Corning) and imaged using the OperaPhenix Live imaging System (PerkinElmer). The imaging of YFP+ cells occurred at 488 nm every 30 min for 4 days. The data was then analyzed using Harmony’s Columbus software with capability to test progeny of the dividing YFP+ cells. Generation information was collected using Columbus’ division tracking. Cells dead after division were defined as cells that start with a “split”, “cosmos”, “merge”, or “border” fate and recorded as “cosmos” at the end of their tracking. Cells that were defined as live after division start with “split”, “cosmos”, “merge”, or “border” and end with “split” or “end” at the end of their tracking. In order to calculate latency to mitosis, cells with “split”-“split” fate were considered. To calculate latency to apoptosis, all cells with “cosmos” end fate were considered. Percentages of different groups of cells were calculated and analyzed.

#### Cell Cycle Flow Cytometry

For *in vivo* cell cycle analysis, 14 dpi WT and Beclin1 nKO mice were injected with EdU four times two hours apart and dentate gyrus was extracted and digested as described above 24 hours after the first injection. After digestion, enzymatic activity was neutralized with media containing DMEM/F12 and 10% Fetal Bovine Serum (FBS; SH3039603; HyClone) and cells were washed with additional DMEM/F12 to remove excess FBS. The cell suspension was transferred in Percoll media, consisting of 19.8% Percoll (17-0891-02; GE Healthcare Life Sciences), 2.2% 10xPBS (Multicell) and the cells were spun down (500 x g, 12.5 minutes, 4°C). The cells were re-suspended in 200μL of DMEM:F12 media and counted with trypan blue. The cells were then fixed in a PFA-containing buffer (420801; BioLegend) for 20 min in dark at RT, permeabilized by spinning in 1x wash buffer (421002; Biolegend) twice for 10 min, and passed through 40 μm cell strainer (08-771-1, Fisher). Beclin1 nKO and WT cells were then stained for 20 min in dark with EdU cocktail prepared as described above. The sections were then washed, and resuspended in DAPI/1xPBS. The cells were then accessed using a Fortessa (Beckman Coulter) flow cytometer at 405 nm and 561 nm lasers for DAPI, and EdU-Cy3, respectively.

For *in vitro* cell cycle assessment of WT and Beclin1 nKO cells, neurosphere cultures were prepared as described above. On day 10-12 of culture, EdU was added to culture media at a concentration of 10mM and incubated at 37°C for 4 hours. Spheres were then dissociated via incubation in TrypLE (12604-013; Gibco) at 37°C followed by trituration. The cells were washed once with phenol-free DMEM-F12 and counted with trypan blue. The cells were then fixed in a PFA-containing buffer (420801; BioLegend) for 20 min in dark at RT, permeabilized by spinning in 1x wash buffer (421002; Biolegend) twice for 10 min, and passed through 40 μm cell strainer (08-771-1; Fisher). Cells were then stained for 20 min in dark with EdU cocktail prepared as described above, washed, and resuspended in DAPI/1xPBS. The cells were then accessed using a Fortessa (Beckman Coulter) flow cytometer at 405 nm and 561 nm lasers for DAPI and EdU-Cy3, respectively.

All visualization and cell counting of flow cytometry data was performed using FlowJo10 and 11 (BD). Values obtained from different experiments using FlowJo were used in t-test with Prism 6.0 (GraphPad).

#### Single-cell Isolation and Library Preparation

Beclin1 WT and nKO mice were deeply anesthetized and dentate gyrus was extracted and digested as described above. After digestion, cells were washed with media containing DMEM/F12 and 10% Fetal Bovine Serum (FBS; SH3039603; HyClone) to neutralize further enzymatic digestion, then washed with additional DMEM/F12 to remove excess FBS. The cells were transferred in Percoll media, consisting of 19.8% Percoll (17-0891-02; GE Healthcare Life Sciences), 2.2% 10xPBS (Multicell) and spun down (500 x g, 12.5 minutes, 4°C). The cells were re-suspended in 200μL of DMEM:F12 media with 7AAD at a concentration of 50uL/10^6^ cells and transferred to the cell sorter on ice. Approximately 60k WT and 40k Beclin1-null YFP+ cells negative for dead marker 7AAD were sorted using a MoFlo Astriosis cell sorter (Beckman Coulter) into DMEM-F12, spun down to concentrate the pellet, resuspended and counted on a Countess Cell Counter. Cells were sequenced by 10x genomics using Chromium’s Single Cell 3’ v2 chemistry kit. The cDNA libraries were purified, quantified, and then sequenced on the next generation sequencing Illumina NextSeq 500 platform.

#### Cell Clustering, Visualization, Differential Gene Expression and Go Term Analyses

Prior to data analysis, low quality cells were stringently filtered out based on percentage of mitochondrial genes (<6%), detected RNA counts (500-10,000), and detected RNA features (i.e., genes, 500-3,500). Addition, cells with a high microglial signature of Fcer1g (<0.01) and Csf1r (<0.001) gene expression were also filtered out. For unsupervised data clustering we used Satija’s R package Seurat v4.0 ^92^. WT and Beclin1 nKO samples were merged, split, and SCTransformed. Integration features and anchors were then calculated, and samples were combined for an integrated analysis. Principle component analysis (PCA) was performed and the first 10 PCs were used for Uniform Manifold Approximation and Projection (UMAP) and cluster finding at a clustering resolution of 0.8. After the removal of pericytes, oligodendrocytes, and endothelial cells principal components were calculated again, and reduction was performed on the first 20 components and clustered at a resolution of 0.8. For later analyses on the proliferating clusters, similar steps were performed but increasing the number of PCAs to 30 and clustering resolution to 1.2. Differentially expressed genes (DEGs) were found for the proliferating clusters (adjusted p value < 0.05 and more than 1.5-fold change or p value < 0.05) and the full list is shown in supplementary Table S2.

For GO Term analysis of gross lists of DEGs, DEGs were split into upregulated and downregulated first. The lists were then fed to an online Metascape (metascape.org) for mouse species and resulting GO Terms with LogP>4 are shown in ?.

#### Analysis of Cell Trajectories and Lineage-driving Transcriptional Regulators

For Pseudotime analysis, Trapnell’s Monocle^93^ R package was used to pseudotemporally order the cycling clusters, which included aNSCs, S-phase NSCs and NPCs, mitotic cells and cells exiting the cell cycle (CCE1-3). Monocle’s differential gene test was also used to confirm DEGs obtained using Seurat. The cells were classified based on expression of quiescence gene Aldoc (≥1.5), proliferation genes Mki67 (>1) and Dynll1 (≥10), and differentiation-related gene Tubb3 (≥5). Pseudotemporal metadata was then transferred to the original Seurat object to maintain consistent clustering of cycling cell populations. To order cells along the continuum with calculated GO Term scores for different pathways, first gene sets were obtained from the Mouse Gene Informatics (MGI, informatics.jax.org) using the Gene Ontology browser. GO Term scores were then calculated and added to Seurat metadata, and used along with the pseudotime continuum to determine the timing of activation/expression within the cycling cells.

In order to identify fate trajectories of proliferating neural stem and progenitor cells, single cell RNA velocities were calculated for WT and Beclin1 nKO samples using Bergen’s scVelo ^94^. Briefly, seurat object containing cycling cells was split into WT and Beclin1 nKO and each processed separately in scVelo using identical code. Spliced and unspliced counts were used to determine RNA velocity and construct cell fate vectors using a dynamical velocity model. Latent time of cell processes was quantified per each cell cluster allowing to determine 100 cluster-specific top-likelihood driver genes that were obtained for WT and Beclin1 nKO. The resulting gene sets were compared to determine the number of same and unique genes regulating WT and Beclin1 nKO cell cycle progression and exit.

#### Comet Assay

For assessment of DNA damage, dissociated DG cells were sorted from WT and Beclin1 nKO mice at 14 dpi using a Beckman MoFlo Astriosis (Beckman Coulter Canada) for YFP positivity (488-526 nm) and for 7-AAD negativity (571-640nm). Alkaline comet assay was performed according to previously published methods with modifications ^74^. Briefly, frosted slides were pre-coated with normal melting point agarose (1% in 1xTBE), dried, and stored at 4°C. 1x TBE (Tris, borate, and EDTA buffer) was cooled before use and kept on ice. Alkaline electrophoresis buffer (300 mM NaOH, 1 mM EDTA, pH>13) was made fresh and cooled before use. On the day of comet assay, low melting point agarose was dissolved in 1xTBE to a final of 1% and kept in 37°C water bath until use. Sorted cells at a concentration of 10^6^ were combined with low melting point agarose as a ratio of 1:10 and applied to pre-coated slides. Cold lysis solution (R&D) was applied to slides and left at 4°C overnight. Slides were washed with 1x TBE and placed into cold electrophoresis buffer for 30 min to unwind DNA. The current was then applied to slides for 30 min at 1.87 V/cm on ice. Slides were washed and stained with Sybr Gold (S11494; Invitrogen) in 1X PBS for 20 min, partially dried, coverslipped with ImmuMount, and stored flat at 4°C until imaging. The slides were imaged at 20x on an Olympus BX51 epifluorescent microscope at 561 nm. For comet tail analysis, profile analysis was used in OpenComet plugin in ImageJ with no background correction. Extremely dim cells and cells with percent of DNA in tail higher than 70% were excluded from the analysis.

### QUANTIFICATION AND STATISTICAL ANALYSIS

All outcomes are reported as mean ± standard error of the mean (SEM) and were calculated and statistically analyzed using Prism 6.0 (GraphPad). Experiments with two groups were analyzed by a two-tailed student’s t-test. Statistical analysis of three or more groups was performed using an ANOVA test, followed by a Bonferroni post hoc. Statistical significance was defined as *p < 0.05*.

### DATA AND SOFTWARE AVAILABILITY

All the scRNA-seq data have been deposited in the NCBI Gene Expression Omnibus (GEO) under accession number GEO: GSE247493.

## Supplemental Information

**Figure S1. Retroviral assessment of autolysosomes of Beclin1 nKO mice shows reduced autophagy.** (A) Representative images of mCherry+ autolysosomes (red) in WT and Beclin1-null YFP+ (green) retrovirally infected cells. (G) Beclin1 removal is associated with a reduced total number of mCherry+ puncta per cell (mean ± SEM, unpaired t-test). Scale bar represents 5 uM (A); ** p≤0.01.

**Figure S2. Quantification of quiescent NSCs in WT and Beclin1 nKO mice shows no change.** (A) Co-labeling of AldoC (violet) and Mcm2 (blue) with YFP+ (green) recombined WT and Beclin1 nKO in dentate gyrus at 14 dpi shows no change in quiescent NSCs positive for Aldoc and negative for Mcm2 in slices. (B) Removal of Beclin1 resulted in no change in the percentage of YFP+Aldoc+Mcm2-qNSCs at 14 or 30 dpi (mean ± SEM, 2-way ANOVA). Scale bar represents 10 uM.

**Figure S3. Virus-mediated removal of Beclin1 from dentate gyrus NPCs in WT and Beclin1 fl/fl mice has minor effects.** (A) Representative images of GFP+ (green) and RFP+ (red) cells virally infected in the dentate gyrus of WT and Beclin1 fl/fl mice at 14, 30, and 60 dpi. (B) Removal of Beclin1 from dentate gyrus NPCs resulted in an overall modest effect of phenotype on survival of GFP+RFP+ NPCs and their progeny over time (mean ± SEM, 2-way ANOVA). (C) Representative fluorescent images of virally infected WT and Beclin1 fl/fl cells (red and green) at 14 dpi (left) and close-up images of dendritic spines (white) analyzed using Sholl analysis. (D) Sholl analysis results show no difference in the number of spines per 10 uM in WT and Beclin1 fl/fl infected cells (mean ± SEM, unpaired t-test). Scale bars represent 20 uM (A), 10 uM (C left), 5 uM (C right); * p≤0.05.

**Figure S4. Beclin1 loss results in overall reduction of DG immature neurons.** (A) Representative DAB-DCX (black) images of WT and Beclin1 nKO mice at 14, 30, and 60 dpi. (B) Quantification of total DAB-DCX+ cells shows a decrease in Beclin1 nKO mice at 30 and 60 dpi (mean ± SEM, 2-way ANOVA). Scale bar represents 500 uM; ** p≤0.01, *** p≤0.001.

**Figure S5. PCR primers.** (A) Primers used for the genotyping of WT and Beclin1 nKO mice.

**Table S1.** Velocity driver genes ranked in each cluster.

**Table S2.** List of genes differentially expressed in Beclin1-nul NSPCs.

## References

1. Menon, M.B., and Dhamija, S. (2018). Beclin 1 Phosphorylation – at the Center of Autophagy Regulation. Front. Cell Dev. Biol. 6.

2. Kang, R., Zeh, H.J., Lotze, M.T., and Tang, D. (2011). The Beclin 1 network regulates autophagy and apoptosis. Cell Death Differ. 18, 571–580. 10.1038/cdd.2010.191.

3. Funderburk., S.F., Wang, Q.J., and Yue, Z. (2010). Beclin 1-VPS34 complex – At the Crossroads of Autophagy and Beyond. Trends Cell Biol. 20, 355–362. 10.1016/j.tcb.2010.03.002.

4. Noguchi, S., Honda, S., Saitoh, T., Matsumura, H., Nishimura, E., Akira, S., and Shimizu, S. (2019). Beclin 1 regulates recycling endosome and is required for skin development in mice. Commun. Biol. 2, 37. 10.1038/s42003-018-0279-0.

5. Konishi, A., Arakawa, S., Yue, Z., and Shimizu, S. (2012). Involvement of Beclin 1 in engulfment of apoptotic cells. J. Biol. Chem. 287, 13919–13929. 10.1074/jbc.M112.348375.

6. Lemus Silva, E.G., Delgadillo, Y., White, R.E., and Lucin, K.M. (2023). Beclin 1 regulates astrocyte phagocytosis and phagosomal recruitment of retromer. Tissue Cell 82, 102100. 10.1016/j.tice.2023.102100.

7. Hamurcu, Z., Delibaşı, N., Geçene, S., Şener, E.F., Dönmez-Altuntaş, H., Özkul, Y., Canatan, H., and Ozpolat, B. (2018). Targeting LC3 and Beclin-1 autophagy genes suppresses proliferation, survival, migration and invasion by inhibition of Cyclin-D1 and uPAR/Integrin β1/ Src signaling in triple negative breast cancer cells. J. Cancer Res. Clin. Oncol. 144, 415–430. 10.1007/s00432-017-2557-5.

8. Park, J.M., Tougeron, D., Huang, S., Okamoto, K., and Sinicrope, F.A. (2014). Beclin 1 and UVRAG Confer Protection from Radiation-Induced DNA Damage and Maintain Centrosome Stability in Colorectal Cancer Cells. PLOS ONE 9, e100819. 10.1371/journal.pone.0100819.

9. Xu, F., Fang, Y., Yan, L., Xu, L., Zhang, S., Cao, Y., Xu, L., Zhang, X., Xie, J., Jiang, G., et al. (2017). Nuclear localization of Beclin 1 promotes radiation-induced DNA damage repair independent of autophagy. Sci. Rep. 7, 45385. 10.1038/srep45385.

10. Frémont, S., Gérard, A., Galloux, M., Janvier, K., Karess, R.E., and Berlioz-Torrent, C. (2013). Beclin-1 is required for chromosome congression and proper outer kinetochore assembly. EMBO Rep. 14, 364–372. 10.1038/embor.2013.23.

11. Yue, Z., Jin, S., Yang, C., Levine, A.J., and Heintz, N. (2003). Beclin 1, an autophagy gene essential for early embryonic development, is a haploinsufficient tumor suppressor. Proc. Natl. Acad. Sci. U. S. A. 100, 15077–15082. 10.1073/pnas.2436255100.

12. Yazdankhah, M., Farioli-Vecchioli, S., Tonchev, A.B., Stoykova, A., and Cecconi, F. (2014). The autophagy regulators Ambra1 and Beclin 1 are required for adult neurogenesis in the brain subventricular zone. Cell Death Dis. 5, e1403–e1403. 10.1038/cddis.2014.358.

13. McKnight, N.C., Zhong, Y., Wold, M.S., Gong, S., Phillips, G.R., Dou, Z., Zhao, Y., Heintz, N., Zong, W.X., and Yue, Z. (2014). Beclin 1 Is Required for Neuron Viability and Regulates Endosome Pathways via the UVRAG-VPS34 Complex. PLoS Genet. 10, 1–18. 10.1371/journal.pgen.1004626.

14. Xi, Y., Dhaliwal, J.S., Ceizar, M., Vaculik, M., Kumar, K.L., and Lagace, D.C. (2016). Knockout of Atg5 delays the maturation and reduces the survival of adult-generated neurons in the hippocampus. Cell Death Dis. 7, e2127. 10.1038/cddis.2015.406.

15. Furutachi, S., Miya, H., Watanabe, T., Kawai, H., Yamasaki, N., Harada, Y., Imayoshi, I., Nelson, M., Nakayama, K.I., Hirabayashi, Y., et al. (2015). Slowly dividing neural progenitors are an embryonic origin of adult neural stem cells. Nat. Neurosci. 18, 657–665. 10.1038/nn.3989.

16. Maslov, A.Y., Barone, T.A., Plunkett, R.J., and Pruitt, S.C. (2004). Neural Stem Cell Detection, Characterization, and Age-Related Changes in the Subventricular Zone of Mice. J. Neurosci. 24, 1726–1733. 10.1523/JNEUROSCI.4608-03.2004.

17. Chaker, Z., Codega, P., and Doetsch, F. (2016). A mosaic world: puzzles revealed by adult neural stem cell heterogeneity. WIREs Dev. Biol. 5, 640–658. 10.1002/wdev.248.

18. Dulken, B.W., Leeman, D.S., Boutet, S.C., Hebestreit, K., and Brunet, A. (2017). Single-Cell Transcriptomic Analysis Defines Heterogeneity and Transcriptional Dynamics in the Adult Neural Stem Cell Lineage. Cell Rep. 18, 777–790. 10.1016/j.celrep.2016.12.060.

19. Su, Y.-T., Lau, S.-F., Ip, J.P.K., Cheung, K., Cheung, T.H.T., Fu, A.K.Y., and Ip, N.Y. (2019). α2-Chimaerin is essential for neural stem cell homeostasis in mouse adult neurogenesis. Proc. Natl. Acad. Sci. 116, 13651–13660. 10.1073/pnas.1903891116.

20. Ellis, P., Fagan, B.M., Magness, S.T., Hutton, S., Taranova, O., Hayashi, S., McMahon, A., Rao, M., and Pevny, L. (2004). SOX2, a Persistent Marker for Multipotential Neural Stem Cells Derived from Embryonic Stem Cells, the Embryo or the Adult. Dev. Neurosci. 26, 148–165. 10.1159/000082134.

21. Shimozaki, K. (2014). Sox2 transcription network acts as a molecular switch to regulate properties of neural stem cells. World J. Stem Cells 6, 485–490. 10.4252/wjsc.v6.i4.485.

22. Brazel, C.Y., Limke, T.L., Osborne, J.K., Miura, T., Cai, J., Pevny, L., and Rao, M.S. (2005). Sox2 expression defines a heterogeneous population of neurosphere-forming cells in the adult murine brain. Aging Cell 4, 197–207. 10.1111/j.1474-9726.2005.00158.x.

23. Kim, J.S., Park, S.W., Hwang, I., Kim, Y.W., Kim, J.H., and Kim, J.H. (2015). Expression of nestin on endothelial cells and pericytes during vascular development in mouse retina. Invest. Ophthalmol. Vis. Sci. 56, 3403–3403.

24. Almazán, G., Vela, J.M., Molina-Holgado, E., and Guaza, C. (2001). Re-evaluation of nestin as a marker of oligodendrocyte lineage cells. Microsc. Res. Tech. 52, 753–765. 10.1002/jemt.1060.

25. Shin, J., Berg, D.A., Zhu, Y., Shin, J.Y., Song, J., Bonaguidi, M.A., Enikolopov, G., Nauen, D.W., Christian, K.M., Ming, G., et al. (2015). Single-Cell RNA-Seq with Waterfall Reveals Molecular Cascades underlying Adult Neurogenesis. Cell Stem Cell 17, 360–372. 10.1016/j.stem.2015.07.013.

26. Basak, O., Krieger, T.G., Muraro, M.J., Wiebrands, K., Stange, D.E., Frias-Aldeguer, J., Rivron, N.C., Wetering, M. van de, Es, J.H. van, Oudenaarden, A. van, et al. (2018). Troy+ brain stem cells cycle through quiescence and regulate their number by sensing niche occupancy. Proc. Natl. Acad. Sci. 115, E610–E619. 10.1073/pnas.1715911114.

27. Zerjatke, T., Gak, I.A., Kirova, D., Fuhrmann, M., Daniel, K., Gonciarz, M., Müller, D., Glauche, I., and Mansfeld, J. (2017). Quantitative Cell Cycle Analysis Based on an Endogenous All-in-One Reporter for Cell Tracking and Classification. Cell Rep. 19, 1953–1966. 10.1016/j.celrep.2017.05.022.

28. Pereira, P.D., Serra-Caetano, A., Cabrita, M., Bekman, E., Braga, J., Rino, J., Santus, R., Filipe, P.L., Sousa, A.E., and Ferreira, J.A. (2017). Quantification of cell cycle kinetics by EdU (5-ethynyl-2′-deoxyuridine)-coupled-fluorescence-intensity analysis. Oncotarget 8, 40514–40532. 10.18632/oncotarget.17121.

29. Yang, C.-P., Gilley, J.A., Zhang, G., and Kernie, S.G. (2011). ApoE is required for maintenance of the dentate gyrus neural progenitor pool. Dev. Camb. Engl. 138, 4351–4362. 10.1242/dev.065540.

30. Satija, R., Farrell, J.A., Gennert, D., Schier, A.F., and Regev, A. (2015). Spatial reconstruction of single-cell gene expression data. Nat. Biotechnol. 33, 495–502. 10.1038/nbt.3192.

31. Bergen, V., Lange, M., Peidli, S., Wolf, F.A., and Theis, F.J. (2019). Generalizing RNA velocity to transient cell states through dynamical modeling. bioRxiv, 820936. 10.1101/820936.

32. Jiang, Y., Bruin, A. de, Caldas, H., Fangusaro, J., Hayes, J., Conway, E.M., Robinson, M.L., and Altura, R.A. (2005). Essential Role for Survivin in Early Brain Development. J. Neurosci. 25, 6962–6970. 10.1523/JNEUROSCI.1446-05.2005.

33. Wiedemuth, R., Klink, B., Töpfer, K., Schröck, E., Schackert, G., Tatsuka, M., and Temme, A. (2014). Survivin safeguards chromosome numbers and protects from aneuploidy independently from p53. Mol. Cancer 13, 107. 10.1186/1476-4598-13-107.

34. Piper, M., Barry, G., Harvey, T., Mcleay, R., Smith, A., Harris, L., Mason, S., Stringer, B., Day, B., Wray, N., et al. (2014). NFIB-Mediated Repression of the Epigenetic Factor Ezh2 Regulates Cortical Development. J. Neurosci. Off. J. Soc. Neurosci. 34, 2921–2930. 10.1523/JNEUROSCI.2319-13.2014.

35. Clark, B.S., Stein-O’Brien, G.L., Shiau, F., Cannon, G.H., Davis-Marcisak, E., Sherman, T., Santiago, C.P., Hoang, T.V., Rajaii, F., James-Esposito, R.E., et al. (2019). Single-Cell RNA-Seq Analysis of Retinal Development Identifies NFI Factors as Regulating Mitotic Exit and Late-Born Cell Specification. Neuron 102, 1111–1126.e5. 10.1016/j.neuron.2019.04.010.

36. Regnell, C.E., Hildrestrand, G.A., Sejersted, Y., Medin, T., Moldestad, O., Rolseth, V., Krokeide, S.Z., Suganthan, R., Luna, L., Bjørås, M., et al. (2012). Hippocampal adult neurogenesis is maintained by Neil3-dependent repair of oxidative DNA lesions in neural progenitor cells. Cell Rep. 2, 503–510. 10.1016/j.celrep.2012.08.008.

37. Paonessa, F., Evans, L.D., Solanki, R., Larrieu, D., Wray, S., Hardy, J., Jackson, S.P., and Livesey, F.J. (2019). Microtubules Deform the Nuclear Membrane and Disrupt Nucleocytoplasmic Transport in Tau-Mediated Frontotemporal Dementia. Cell Rep. 26, 582–593.e5. 10.1016/j.celrep.2018.12.085.

38. Leveille, E., Ross, O.A., and Gan-Or, Z. (2021). Tau and MAPT genetics in tauopathies and synucleinopathies. Parkinsonism Relat. Disord. 90, 142–154. 10.1016/j.parkreldis.2021.09.008.

39. Verdaasdonk, J.S., and Bloom, K. (2011). Centromeres: unique chromatin structures that drive chromosome segregation. Nat. Rev. Mol. Cell Biol. 12, 320–333.

40. Pesenti, M.E., Raisch, T., Conti, D., Walstein, K., Hoffmann, I., Vogt, D., Prumbaum, D., Vetter, I.R., Raunser, S., and Musacchio, A. (2022). Structure of the human inner kinetochore CCAN complex and its significance for human centromere organization. Mol. Cell 82, 2113–2131.e8. 10.1016/j.molcel.2022.04.027.

41. Waitzman, J.S., and Rice, S.E. (2014). Mechanism and regulation of kinesin-5, an essential motor for the mitotic spindle. Biol. Cell 106, 1–12. 10.1111/boc.201300054.

42. Pryzhkova, M.V., and Jordan, P.W. (2016). Conditional mutation of Smc5 in mouse embryonic stem cells perturbs condensin localization and mitotic progression. J. Cell Sci. 129, 1619–1634. 10.1242/jcs.179036.

43. Cross, R.A., and McAinsh, A. (2014). Prime movers: the mechanochemistry of mitotic kinesins. Nat. Rev. Mol. Cell Biol. 15, 257–271. 10.1038/nrm3768.

44. Paul, M.R., Hochwagen, A., and Ercan, S. (2019). Condensin action and compaction. Curr. Genet. 65, 407–415. 10.1007/s00294-018-0899-4.

45. Martin, C.-A., Murray, J.E., Carroll, P., Leitch, A., Mackenzie, K.J., Halachev, M., Fetit, A.E., Keith, C., Bicknell, L.S., Fluteau, A., et al. (2016). Mutations in genes encoding condensin complex proteins cause microcephaly through decatenation failure at mitosis. Genes Dev. 30, 2158–2172. 10.1101/gad.286351.116.

46. Barra, V., and Fachinetti, D. (2018). The dark side of centromeres: types, causes and consequences of structural abnormalities implicating centromeric DNA. Nat. Commun. 9, 4340. 10.1038/s41467-018-06545-y.

47. Wu, N., and Yu, H. (2012). The Smc complexes in DNA damage response. Cell Biosci. 2, 5. 10.1186/2045-3701-2-5.

48. Kim, J.M. (2022). Molecular Link between DNA Damage Response and Microtubule Dynamics. Int. J. Mol. Sci. 23, 6986. 10.3390/ijms23136986.

49. Thompson, A.F., Blackburn, P.R., Arons, N.S., Stevens, S.N., Babovic-Vuksanovic, D., Lian, J.B., Klee, E.W., and Stumpff, J. (2022). Pathogenic mutations in the chromokinesin KIF22 disrupt anaphase chromosome segregation. eLife 11, e78653. 10.7554/eLife.78653.

50. McCleland, M.L., Kallio, M.J., Barrett-Wilt, G.A., Kestner, C.A., Shabanowitz, J., Hunt, D.F., Gorbsky, G.J., and Stukenberg, P.T. (2004). The vertebrate Ndc80 complex contains Spc24 and Spc25 homologs, which are required to establish and maintain kinetochore-microtubule attachment. Curr. Biol. CB 14, 131–137. 10.1016/j.cub.2003.12.058.

51. Cavazza, T., and Vernos, I. (2016). The RanGTP Pathway: From Nucleo-Cytoplasmic Transport to Spindle Assembly and Beyond. Front. Cell Dev. Biol. 3. 10.3389/fcell.2015.00082.

52. Joseph, J., Liu, S.-T., Jablonski, S.A., Yen, T.J., and Dasso, M. (2004). The RanGAP1-RanBP2 Complex Is Essential for Microtubule-Kinetochore Interactions In Vivo. Curr. Biol. 14, 611–617. 10.1016/j.cub.2004.03.031.

53. Arnaoutov, A., and Dasso, M. (2003). The Ran GTPase Regulates Kinetochore Function. Dev. Cell 5, 99–111. 10.1016/S1534-5807(03)00194-1.

54. D’Angiolella, V., Donato, V., Forrester, F.M., Jeong, Y.-T., Pellacani, C., Kudo, Y., Saraf, A., Florens, L., Washburn, M.P., and Pagano, M. (2012). The Cyclin F-Ribonucleotide Reductase M2 axis controls genome integrity and DNA repair. Cell 149, 1023–1034. 10.1016/j.cell.2012.03.043.

55. Boudhraa, Z., Carmona, E., Provencher, D., and Mes-Masson, A.-M. (2020). Ran GTPase: A Key Player in Tumor Progression and Metastasis. Front. Cell Dev. Biol. 8.

56. Pommier, Y., Nussenzweig, A., Takeda, S., and Austin, C. (2022). Human topoisomerases and their roles in genome stability and organization. Nat. Rev. Mol. Cell Biol. 23, 407–427. 10.1038/s41580-022-00452-3.

57. De Biasio, A., de Opakua, A.I., Mortuza, G.B., Molina, R., Cordeiro, T.N., Castillo, F., Villate, M., Merino, N., Delgado, S., Gil-Cartón, D., et al. (2015). Structure of p15(PAF)-PCNA complex and implications for clamp sliding during DNA replication and repair. Nat. Commun. 6, 6439. 10.1038/ncomms7439.

58. Bolcun-Filas, E., Costa, Y., Speed, R., Taggart, M., Benavente, R., De Rooij, D.G., and Cooke, H.J. (2007). SYCE2 is required for synaptonemal complex assembly, double strand break repair, and homologous recombination. J. Cell Biol. 176, 741–747. 10.1083/jcb.200610027.

59. Morimoto, Y., Tokumitsu, A., Sone, T., Hirota, Y., Tamura, R., Sakamoto, A., Nakajima, K., Toda, M., Kawakami, Y., Okano, H., et al. (2022). TPT1 Supports Proliferation of Neural Stem/Progenitor Cells and Brain Tumor Initiating Cells Regulated by Macrophage Migration Inhibitory Factor (MIF). Neurochem. Res. 47, 2741–2756. 10.1007/s11064-022-03629-6.

60. Gonzalez-Franquesa, A., Stocks, B., Borg, M.L., Kuefner, M., Dalbram, E., Nielsen, T.S., Agrawal, A., Pankratova, S., Chibalin, A.V., Karlsson, H.K.R., et al. (2021). Discovery of thymosin β4 as a human exerkine and growth factor. Am. J. Physiol. Cell Physiol. 321, C770– C778. 10.1152/ajpcell.00263.2021.

61. Cai, Q., Lu, L., Tian, J.-H., Zhu, Y.-B., Qiao, H., and Sheng, Z.-H. (2010). Snapin-regulated late endosomal transport is critical for efficient autophagy-lysosomal function in neurons. Neuron 68, 73–86. 10.1016/j.neuron.2010.09.022.

62. Beesley, P.W., Herrera-Molina, R., Smalla, K.-H., and Seidenbecher, C. (2014). The Neuroplastin adhesion molecules: key regulators of neuronal plasticity and synaptic function. J. Neurochem. 131, 268–283. 10.1111/jnc.12816.

63. Mesman, S., Bakker, R., and Smidt, M.P. (2020). Tcf4 is required for correct brain development during embryogenesis. Mol. Cell. Neurosci. 106, 103502. 10.1016/j.mcn.2020.103502.

64. Trimbuch, T., and Rosenmund, C. (2016). Should I stop or should I go? The role of complexin in neurotransmitter release. Nat. Rev. Neurosci. 17, 118–125. 10.1038/nrn.2015.16.

65. Willems, P.H.G.M., Rossignol, R., Dieteren, C.E.J., Murphy, M.P., and Koopman, W.J.H. (2015). Redox Homeostasis and Mitochondrial Dynamics. Cell Metab. 22, 207–218. 10.1016/j.cmet.2015.06.006.

66. Forsha, D., Church, C., Wazny, P., and Poyton, R.O. (2001). Structure and function of Pet100p, a molecular chaperone required for the assembly of cytochrome c oxidase in Saccharomyces cerevisiae. Biochem. Soc. Trans. 29, 436–441. 10.1042/bst0290436.

67. Suhane, S., Kanzaki, H., Arumugaswami, V., Murali, R., and Ramanujan, V.K. (2013). Mitochondrial NDUFS3 regulates the ROS-mediated onset of metabolic switch in transformed cells. Biol. Open 2, 295–305. 10.1242/bio.20133244.

68. Chen, W., Wang, H., Tao, S., Zheng, Y., Wu, W., Lian, F., Jaramillo, M., Fang, D., and Zhang, D.D. (2013). Tumor protein translationally controlled 1 is a p53 target gene that promotes cell survival. Cell Cycle 12, 2321–2328. 10.4161/cc.25404.

69. Zhang, J., de Toledo, S.M., Pandey, B.N., Guo, G., Pain, D., Li, H., and Azzam, E.I. (2012). Role of the translationally controlled tumor protein in DNA damage sensing and repair. Proc. Natl. Acad. Sci. U. S. A. 109, E926–933. 10.1073/pnas.1106300109.

70. de Moraes, E., Dar, N.A., de Moura Gallo, C.V., and Hainaut, P. (2007). Cross-talks between cyclooxygenase-2 and tumor suppressor protein p53: Balancing life and death during inflammatory stress and carcinogenesis. Int. J. Cancer 121, 929–937. 10.1002/ijc.22899.

71. Budanov, A.V., and Karin, M. (2008). p53 Target Genes Sestrin1 and Sestrin2 Connect Genotoxic Stress and mTOR Signaling. Cell 134, 451–460. 10.1016/j.cell.2008.06.028.

72. Levine, M.S., and Holland, A.J. (2018). The impact of mitotic errors on cell proliferation and tumorigenesis. Genes Dev. 32, 620–638. 10.1101/gad.314351.118.

73. Hayashi, M.T., and Karlseder, J. (2013). DNA damage associated with mitosis and cytokinesis failure. Oncogene 32, 4593–4601. 10.1038/onc.2012.615.

74. Nowsheen, S., Xia, F., and Yang, E.S. (2012). Assaying DNA Damage in Hippocampal Neurons Using the Comet Assay. J. Vis. Exp., 50049. 10.3791/50049.

75. Wang, C., Haas, M., Yeo, S.K., Sebti, S., Fernández, Á.F., Zou, Z., Levine, B., and Guan, J.-L. (2021). Enhanced autophagy in Becn1F121A/F121A knockin mice counteracts aging-related neural stem cell exhaustion and dysfunction. Autophagy 0, 1–14. 10.1080/15548627.2021.1936358.

76. You, S.Y., Park, Y.S., Jeon, H.-J., Cho, D.-H., Jeon, H.B., Kim, S.H., Chang, J.W., Kim, J.-S., and Oh, J.S. (2016). Beclin-1 knockdown shows abscission failure but not autophagy defect during oocyte meiotic maturation. Cell Cycle 15, 1611–1619. 10.1080/15384101.2016.1181235.

77. Mathew, R., Kongara, S., Beaudoin, B., Karp, C.M., Bray, K., Degenhardt, K., Chen, G., Jin, S., and White, E. (2007). Autophagy suppresses tumor progression by limiting chromosomal instability. Genes Dev. 21, 1367–1381. 10.1101/gad.1545107.

78. Karantza-Wadsworth, V., Patel, S., Kravchuk, O., Chen, G., Mathew, R., Jin, S., and White, E. (2007). Autophagy mitigates metabolic stress and genome damage in mammary tumorigenesis. Genes Dev. 21, 1621–1635. 10.1101/gad.1565707.

79. Delaney, J.R., Patel, C.B., Bapat, J., Jones, C.M., Ramos-Zapatero, M., Ortell, K.K., Tanios, R., Haghighiabyaneh, M., Axelrod, J., DeStefano, J.W., et al. (2020). Autophagy gene haploinsufficiency drives chromosome instability, increases migration, and promotes early ovarian tumors. PLoS Genet. 16, e1008558. 10.1371/journal.pgen.1008558.

80. Tran, S., Fairlie, W.D., and Lee, E.F. (2021). BECLIN1: Protein Structure, Function and Regulation. Cells 10, 1522. 10.3390/cells10061522.

81. Kaur, S., and Changotra, H. (2020). The beclin 1 interactome: Modification and roles in the pathology of autophagy-related disorders. Biochimie 175, 34–49. 10.1016/j.biochi.2020.04.025.

82. Wirawan, E., Lippens, S., Vanden Berghe, T., Romagnoli, A., Fimia, G.M., Piacentini, M., and Vandenabeele, P. (2012). Beclin1: A role in membrane dynamics and beyond. Autophagy 8, 6–17. 10.4161/auto.8.1.16645.

83. Niu, T.-K., Cheng, Y., Ren, X., and Yang, J.-M. (2010). Interaction of Beclin 1 with survivin regulates sensitivity of human glioma cells to TRAIL-induced apoptosis. FEBS Lett. 584, 3519– 3524. 10.1016/j.febslet.2010.07.018.

84. Qu, X., Yu, J., Bhagat, G., Furuya, N., Hibshoosh, H., Troxel, A., Rosen, J., Eskelinen, E.-L., Mizushima, N., Ohsumi, Y., et al. (2003). Promotion of tumorigenesis by heterozygous disruption of the beclin 1 autophagy gene. J. Clin. Invest. 112, 1809–1820. 10.1172/JCI200320039.

85. Dhaliwal, J., Trinkle-Mulcahy, L., and Lagace, D.C. (2017). Autophagy and Adult Neurogenesis: Discoveries Made Half a Century Ago Yet in their Infancy of being Connected. Brain Plast. 3, 99–110. 10.3233/BPL-170047.

86. Wang, C., Chen, S., Yeo, S., Karsli-Uzunbas, G., White, E., Mizushima, N., Virgin, H.W., and Guan, J.-L. (2016). Elevated p62/SQSTM1 determines the fate of autophagy-deficient neural stem cells by increasing superoxide. J. Cell Biol. 212, 545–560. 10.1083/jcb.201507023.

87. Jung, S., Choe, S., Woo, H., Jeong, H., An, H.-K., Moon, H., Ryu, H.Y., Yeo, B.K., Lee, Y.W., Choi, H., et al. (2020). Autophagic death of neural stem cells mediates chronic stress-induced decline of adult hippocampal neurogenesis and cognitive deficits. Autophagy 16, 512–530. 10.1080/15548627.2019.1630222.

88. Schäffner, I., Minakaki, G., Khan, M.A., Balta, E.-A., Schlötzer-Schrehardt, U., Schwarz, T.J., Beckervordersandforth, R., Winner, B., Webb, A.E., DePinho, R.A., et al. (2018). FoxO Function Is Essential for Maintenance of Autophagic Flux and Neuronal Morphogenesis in Adult Neurogenesis. Neuron 99, 1188–1203.e6. 10.1016/j.neuron.2018.08.017.

89. Wang, C., Liang, C.-C., Bian, Z.C., Zhu, Y., and Guan, J.-L. (2013). FIP200 is required for maintenance and differentiation of postnatal neural stem cells. Nat. Neurosci. 16, 532–542. 10.1038/nn.3365.

90. Liu, H., Wang, C., Yi, F., Yeo, S., Haas, M., Tang, X., and Guan, J.-L. (2021). Non-canonical function of FIP200 is required for neural stem cell maintenance and differentiation by limiting TBK1 activation and p62 aggregate formation. Sci. Rep. 11, 23907. 10.1038/s41598-021-03404-7.

91. Wang, C., Yeo, S., Haas, M.A., and Guan, J.-L. (2017). Autophagy gene FIP200 in neural progenitors non–cell autonomously controls differentiation by regulating microglia. J. Cell Biol. 216, 2581–2596. 10.1083/jcb.201609093.

92. Stuart, T., Butler, A., Hoffman, P., Hafemeister, C., Papalexi, E., Mauck, W.M., Hao, Y., Stoeckius, M., Smibert, P., and Satija, R. (2019). Comprehensive Integration of Single-Cell Data. Cell 177, 1888–1902.e21. 10.1016/j.cell.2019.05.031.

93. Trapnell, C., Cacchiarelli, D., Grimsby, J., Pokharel, P., Li, S., Morse, M., Lennon, N.J., Livak, K.J., Mikkelsen, T.S., and Rinn, J.L. (2014). The dynamics and regulators of cell fate decisions are revealed by pseudotemporal ordering of single cells. Nat. Biotechnol. 32, 381–386. 10.1038/nbt.2859.

94. Bergen, V., Lange, M., Peidli, S., Wolf, F.A., and Theis, F.J. (2020). Generalizing RNA velocity to transient cell states through dynamical modeling. Nat. Biotechnol. 38, 1408–1414. 10.1038/s41587-020-0591-3.

95. Imayoshi, I., Ohtsuka, T., Metzger, D., Chambon, P., and Kageyama, R. (2006). Temporal regulation of Cre recombinase activity in neural stem cells. genesis 44, 233–238. 10.1002/dvg.20212.

96. Srinivas, S., Watanabe, T., Lin, C.-S., William, C.M., Tanabe, Y., Jessell, T.M., and Costantini, F. (2001). Cre reporter strains produced by targeted insertion of EYFP and ECFP into the ROSA26 locus. BMC Dev. Biol. 1, 4. 10.1186/1471-213X-1-4.

97. Truett, G.E., Heeger, P., Mynatt, R.L., Truett, A.A., Walker, J.A., and Warman, M.L. (2000). Preparation of PCR-quality mouse genomic DNA with hot sodium hydroxide and tris (HotSHOT). BioTechniques 29, 52, 54. 10.2144/00291bm09.

98. Lagace, D.C., Whitman, M.C., Noonan, M.A., Ables, J.L., DeCarolis, N.A., Arguello, A.A., Donovan, M.H., Fischer, S.J., Farnbauch, L.A., Beech, R.D., et al. (2007). Dynamic Contribution of Nestin-Expressing Stem Cells to Adult Neurogenesis. J. Neurosci. 27, 12623–12629. 10.1523/JNEUROSCI.3812-07.2007.

99. Tashiro, A., Zhao, C., Suh, H., and Gage, F.H. (2015). Purification and Injection of Retroviral Vectors. Cold Spring Harb. Protoc. 2015, pdb.prot086371. 10.1101/pdb.prot086371.

100. Imayoshi, I., Sakamoto, M., Ohtsuka, T., Takao, K., Miyakawa, T., Yamaguchi, M., Mori, K., Ikeda, T., Itohara, S., and Kageyama, R. (2008). Roles of continuous neurogenesis in the structural and functional integrity of the adult forebrain. Nat. Neurosci. 11, 1153–1161. 10.1038/nn.2185.

101. Dhaliwal, J., and Lagace, D.C. (2011). Visualization and genetic manipulation of adult neurogenesis using transgenic mice. Eur. J. Neurosci. 33, 1025–1036. 10.1111/j.1460-9568.2011.07600.x.

102. Lagace, D.C., Donovan, M.H., DeCarolis, N.A., Farnbauch, L.A., Malhotra, S., Berton, O., Nestler, E.J., Krishnan, V., and Eisch, A.J. (2010). Adult hippocampal neurogenesis is functionally important for stress-induced social avoidance. Proc. Natl. Acad. Sci. 107, 4436– 4441. 10.1073/pnas.0910072107.

103. Zhao, C., Teng, E.M., Summers, R.G., Ming, G., and Gage, F.H. (2006). Distinct Morphological Stages of Dentate Granule Neuron Maturation in the Adult Mouse Hippocampus. J. Neurosci. 26, 3–11. 10.1523/JNEUROSCI.3648-05.2006.

104. Walker, T.L., and Kempermann, G. (2014). One Mouse, Two Cultures: Isolation and Culture of Adult Neural Stem Cells from the Two Neurogenic Zones of Individual Mice. J. Vis. Exp. JoVE. 10.3791/51225.

105. Hagihara, H., Toyama, K., Yamasaki, N., and Miyakawa, T. (2009). Dissection of Hippocampal Dentate Gyrus from Adult Mouse. J. Vis. Exp. JoVE, 1543. 10.3791/1543.

106. Babu, H., Claasen, J.-H., Kannan, S., Rünker, A.E., Palmer, T., and Kempermann, G. (2011). A Protocol for Isolation and Enriched Monolayer Cultivation of Neural Precursor Cells from Mouse Dentate Gyrus. Front. Neurosci. 5. 10.3389/fnins.2011.00089.

107. Babu, H., Cheung, G., Kettenmann, H., Palmer, T.D., and Kempermann, G. (2007). Enriched Monolayer Precursor Cell Cultures from Micro-Dissected Adult Mouse Dentate Gyrus Yield Functional Granule Cell-Like Neurons. PLOS ONE 2, e388. 10.1371/journal.pone.0000388.

