## Supplemental Information for "Mitotic Activity and DNA Maintenance of Adult Neural Stem Cells is Regulated by Beclin1"

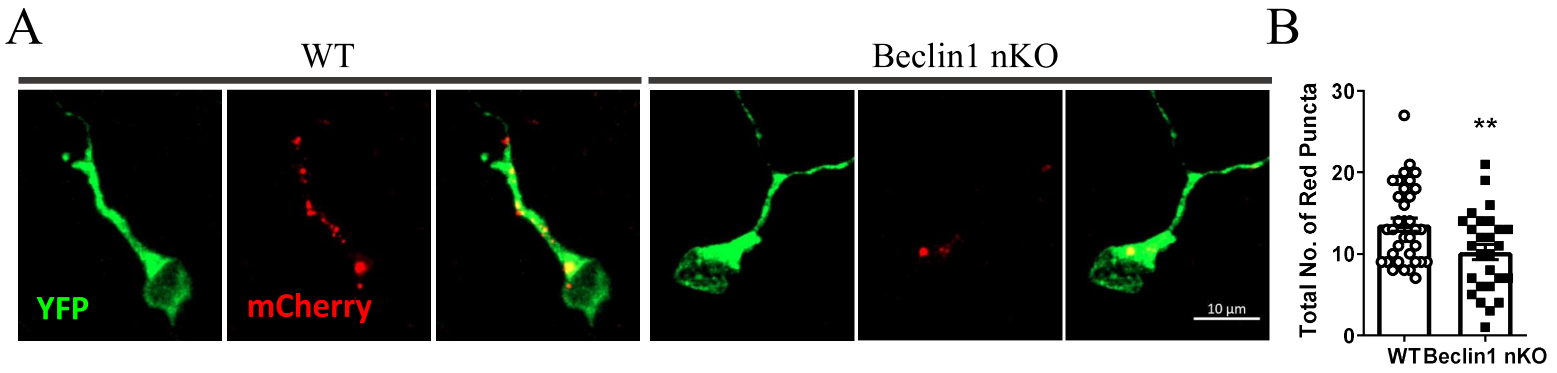


Supplemental Figure 1

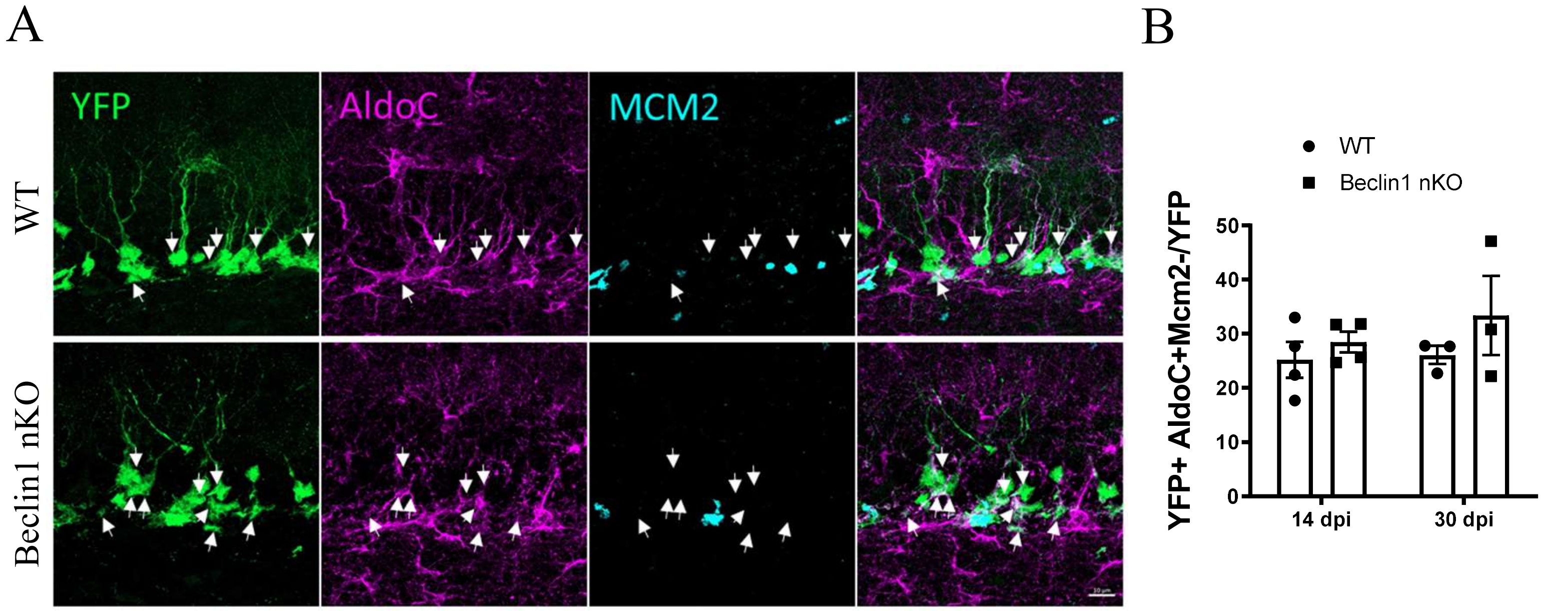


Supplemental Figure 2

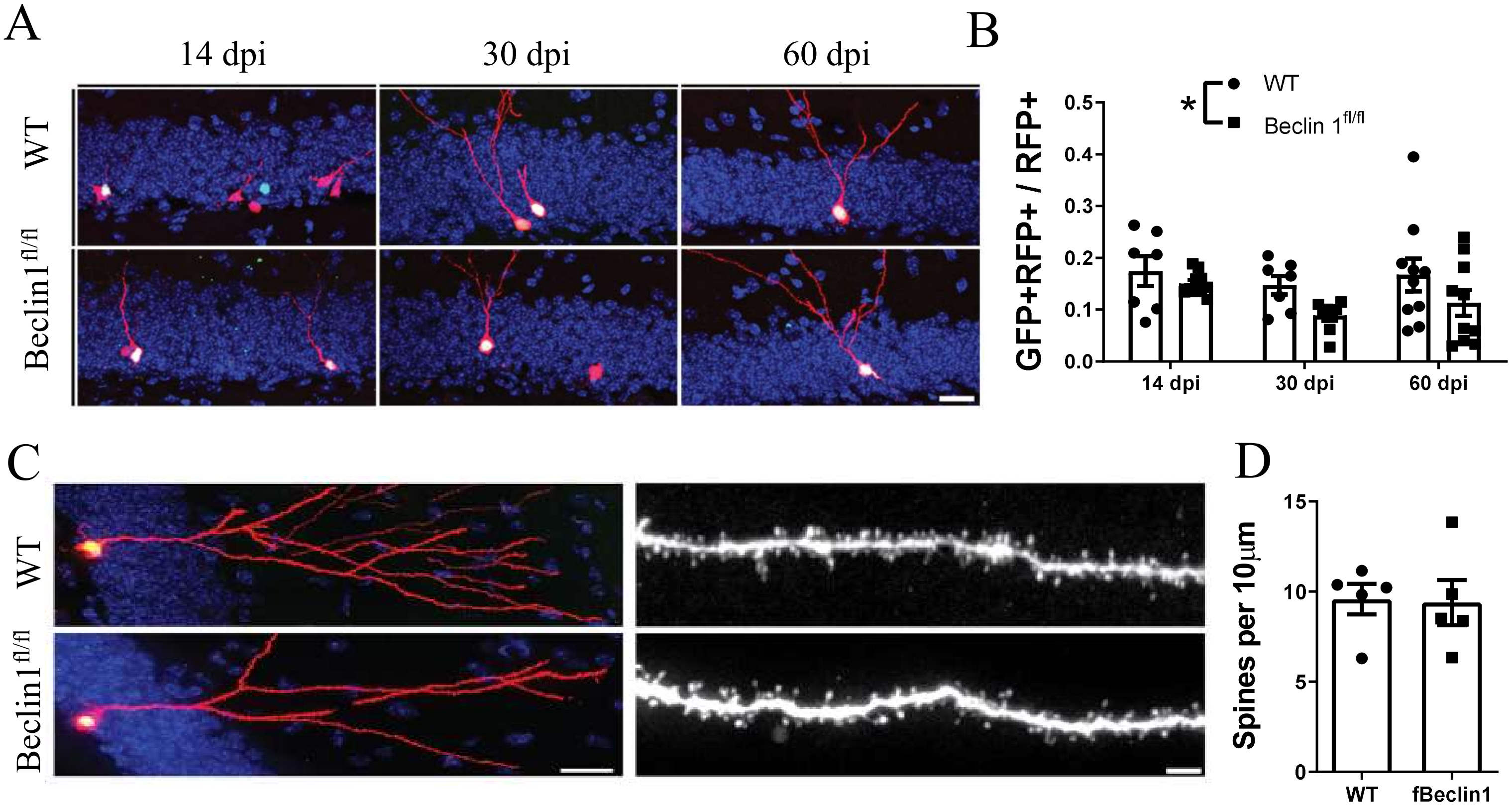


Supplemental Figure 3

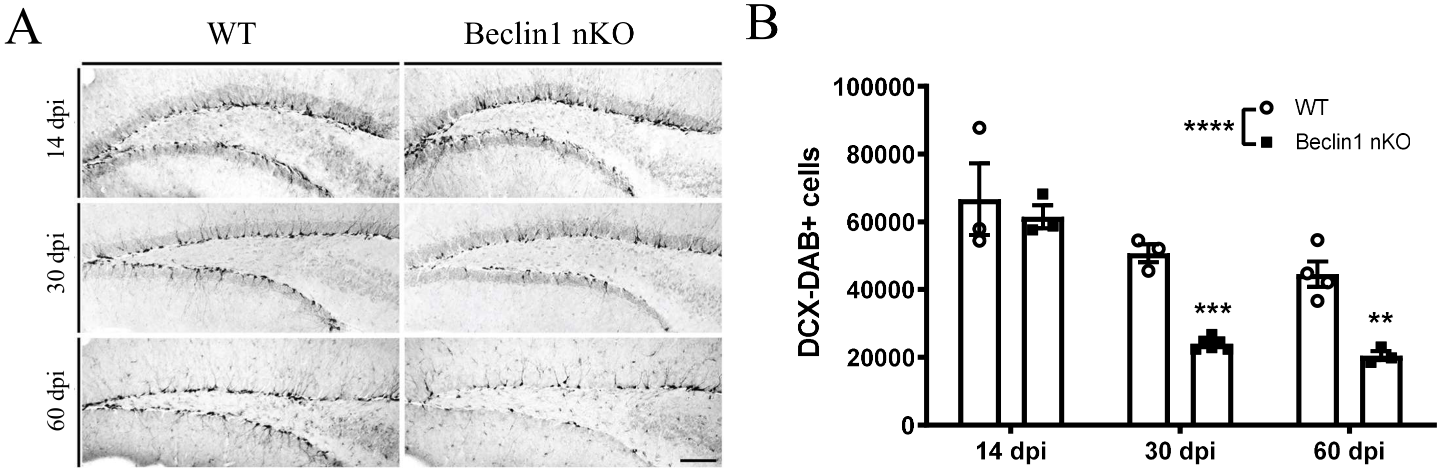


Supplemental Figure 4

A

| Gene | | 5' Primer | 3' Primer | Size (bp) |
|---|---|---|---|---|
| CreER$^{T2}$ | +Control | P26: 5'–CTAGGCCACAGAATTGAAAGATCT–3' | P27: 5'–GTAGGTGGAAATTCTAGCATCATCC–3' | 324 |
| | Transgene | P24: 5'–GCGGTCTGGCAGTAAAAACTATC–3' | P25: 5'–GTGAAACAGCATTGCTGTCACTT–3' | 100 |
| YFP | WT | P21: 5'–GGAGCGGGAGAAATGGATATG–3' | P20: 5'–GCGAAGAGTTTGTCCTCAACC–3' | 560 |
| | Transgene | P21: 5'–GGAGCGGGAGAAATGGATATG–3' | P19: 5'–AAAGTCGCTCTGAGTTGTTAT–3' | 310 |
| fBeclin1 | WT | P70: 5'–CCACCACCAAGGCAGCGGGTAG–3' | P69: 5'–TCACTGATGGCTCTAACCTCAACTCGTC–3' | 650 |
| | Transgene | P70: 5'–CCACCACCAAGGCAGCGGGTAG–3' | P69: 5'–TCACTGATGGCTCTAACCTCAACTCGTC–3' | 850 |

Supplemental Figure 5

| Early M | | Late M | | CCE1 | | CCE2 | | CCE3 | |
|---|---|---|---|---|---|---|---|---|---|
| WT | KO | WT | KO | WT | KO | WT | KO | WT | KO |
| Pclaf | Kntc1 | Cdc25c | Dbx2 | Actn1 | Sstr2 | Klf3 | Nfib | Igfbpl1 | Nfib |
| Pbk | Neil3 | Ptprz1 | Prdx1 | Ptprz1 | Spock2 | Aplp1 | Prdm8 | Nol4 | Nkain4 |
| Birc5 | Shcbp1 | Birc5 | Notch1 | Nfib | Mt1 | Eif4g2 | Mapt | Tubb3 | Sstr2 |
| Bub1b | Spc25 | Ezh2 | Pdlim4 | Stmn1 | Cks2 | Kctd13 | Fam174b | Lzts1 | Kcnb2 |
| Aurka | Ttc28 | Igfbpl1 | Ttc28 | Lzts1 | Dusp15 | 1300017J02Rik | Kcnb2 | Stmn1 | Basp1 |
| Igfbpl1 | Spag5 | H2afy | Neil3 | Nol4 | Fam174b | Igfbpl1 | Dcx | 1300017J02Rik | Cops7a |
| Cdca2 | Mapt | Spc24 | Shcbp1 | Eif4g2 | Cdk1 | Atpif1 | Epha4 | Aplp1 | Fam174b |
| Spc25 | Zmym1 | Basp1 | Nfib | Ezh2 | Fosb | Nol4 | Igfbpl1 | Palm | Mdga1 |
| Lockd | Ska1 | Rgma | Cdk1 | Chchd10 | Haus4 | Elavl4 | Ttc28 | Atpif1 | Polrmt |
| Ttk | Sema3c | Sox9 | Gm10561 | Nucks1 | Pak3 | Fdft1 | Sema3c | Eif4g2 | Sccpdh |
| Ezh2 | Kif4 | Stmn1 | Kntc1 | Elavl4 | Nfib | Nfasc | Nyap1 | Chmp5 | Tfap2c |
| Iqgap3 | Tuba1c | Rad21 | Sox9 | Dek | Cdh24 | Mdga1 | Ap3b2 | Stmn2 | Cdh24 |
| Top2a | Cdk1 | Plcd4 | Mdga1 | Chmp5 | Basp1 | Kif1b | Grk6 | Gm14486 | Kcnh7 |
| Mis18bp1 | Cdkn3 | Ckap2 | Ska1 | Aldh2 | Zmym1 | Tubb3 | Eif3f | Igsf8 | Ttc28 |
| Prr11 | Asf1b | Ncapd2 | Nnat | Cklf | Prc1 | Khdrbs2 | Nrep | Fdft1 | Dcx |
| Cenpe | Mdga1 | Eif4g2 | Basp1 | Pak7 | Akap6 | Srrm4 | Ccdc28b | Hspa14 | 1700025G04Rik |
| Prc1 | Csrp1 | Brd8 | Spag5 | Mdga1 | Nr2e1 | Cttnbp2 | Fam49a | Zfp28 | Mapt |
| Sox9 | Gem | Sfxn5 | Slc1a2 | Palm | Pak7 | Cenpt | Basp1 | Scrt2 | Pak7 |
| Knl1 | Tpcn1 | Smc2 | Cks2 | Cenpt | Polrmt | Fcgrt | Neurod1 | Calm2 | Ggta1 |
| Spc24 | Zfp36l1 | Cenpp | Kif2c | Lhx2 | Serping1 | Stmn2 | Gm17750 | Sh3kbp1 | Myt1l |
| Anln | Bub1b | Gem | Bub1b | Zfp202 | Zfp36l1 | Clybl | Rad18 | Synpr | Prkab1 |
| Kif15 | Sox9 | Aunip | B4galt1 | Ripk1 | Mapt | Frmd5 | Nnat | Cenpt | Prdm8 |
| Nol4 | Prdx1 | Pdik1l | Spc25 | Scrt2 | 1700025G04Rik | Chmp5 | Id2 | Pacsin1 | Epha4 |
| Cklf | Igfbpl1 | Knl1 | Tcf4 | Atpif1 | Atp5b | Zfp532 | Ppp3ca | Nfasc | Snap25 |
| Tpx2 | Cdc25c | Ptn | Myo10 | Insc | Ptprf | Nasp | Satb1 | Cep72 | Nol4 |
| H2afy | Cdca2 | G2e3 | Drd2 | Nasp | Irf9 | Mndal | Mdga1 | Nrn1 | Nrep |
| Trip13 | Bcl11a | Ckap2l | Cttnbp2 | Epha4 | Bcl11a | Stxbp1 | Ppp1r14c | Lemd1 | Tubb3 |
| Ckap2 | Epha4 | 44996 | Cdca2 | Nsrp1 | Hdac4 | Gipc1 | Afap1l2 | Zfp599 | Clvs1 |
| Kif4 | Tcf4 | Gm10561 | Fosb | Gsk3b | Hjurp | Vbp1 | Pak3 | Gm17750 | Nrn1 |
| Smc2 | Kifc1 | Nnat | Kifc1 | Poc5 | Tfap2c | Apba2 | Ddx19a | Ttyh3 | Pak3 |
| Smc4 | Ccdc50 | Eef2kmt | Sccpdh | Ints12 | Ccdc34 | Cenpc1 | Nol4 | Lhx2 | Unc5d |
| Mki67 | Cnih3 | Nedd1 | Asf1b | Rasgef1b | Efhd2 | Tnik | Tspan12 | Rnf123 | Igfbpl1 |
| Cenpq | Csad | Aldh2 | Csrp1 | Tcf4 | Dpysl3 | Palm | Mcrs1 | Ugdh | Junb |

| | | | | | | | | | |
|---|---|---|---|---|---|---|---|---|---|
| Tuba1c | Mtfr2 | Pycard | Cops7a | Tubb3 | Enpp3 | Zfp28 | Kcnh7 | Ptpa | Prmt8 |
| Melk | Cep55 | Vim | Cbs | Dopey2 | Mcm4 | Lmnb1 | Atpif1 | Nnat | Scap |
| Cdk1 | Efhd2 | Mapt | Dcx | Car11 | Mgll | Nrn1 | Chpt1 | Pnpla8 | Ccdc50 |
| Ccnb2 | Notch1 | Chmp5 | Kif4 | Clic1 | Spc24 | Slc25a23 | Ogfod3 | Nptx1 | Sema3c |
| Cenpf | Cenpn | Appl2 | Hdac4 | Tmeff1 | Cdk6 | Gnl2 | Csad | Cables1 | St3gal5 |
| Hjurp | Prmt8 | Lzts1 | Gfap | Epha3 | Afap1l2 | Rasgef1b | Rtn1 | Nsrp1 | Shb |
| Trim59 | Melk | Tmpo | Serping1 | Rgs20 | Gm17750 | Ncam1 | Fat1 | Inpp5b | Zfp324 |
| Kif11 | Kif11 | Igdcc3 | Bcl11a | Myl12b | Ccdc28b | Scrt2 | Rcor2 | Kcnk1 | Nnat |
| Brd8 | Nr2e1 | Cenpc1 | Me3 | G2e3 | Phf19 | Odf2 | Plcd1 | Elavl2 | Chpt1 |
| Basp1 | E2f8 | Ticrr | Ankrd37 | Kif1b | Eml5 | Terf1 | Serping1 | Kif21b | Dixdc1 |
| Kif18a | Pak3 | St3gal4 | Gem | Map2k1 | 1810041L15Rik | Neurod1 | Agpat4 | Srrm4 | Gramd3 |
| Ncapd2 | Szrd1 | Synpr | Nr2e1 | Frmd5 | Dixdc1 | Srgap1 | Chp1 | Cmpk1 | Gsk3b |
| Neil3 | Cks2 | Cmpk1 | Tacc3 | Ranbp9 | Amn1 | Tspan5 | Tmem163 | Prkcb | Rnasel |
| Incenp | Slc1a2 | Adk | Epha4 | Neurod4 | Adgrg3 | Epha4 | Unc5d | Ezh2 | Elmo1 |
| Stmn1 | Mns1 | Gpd1 | B2m | Nsg2 | Kbtbd3 | Slc6a6 | Zfpm2 | Nsg2 | Bcl11a |
| H2afv | Fosb | Pard3b | Nusap1 | Cyp39a1 | Cnih3 | Pacsin1 | 2410004B18Rik | Kif1b | Serping1 |
| Tmpo | Wnt7b | Tnfaip8l1 | Prdm8 | Plch1 | Abcf1 | Cryzl1 | Nek7 | Gsk3b | Mmp15 |
| Hmmr | B4galt1 | Clspn | Nol4l | Tmem159 | Psat1 | Rbms1 | Rps2 | Terf1 | Stmn2 |
| Ccdc18 | Mmd2 | Mad2l1 | Csmd2 | Gpd1 | Pkn2 | Ciapin1 | Junb | Vbp1 | Fam208a |
| Kif5c | Akap6 | 2810459M11Rik | Ccdc50 | Igsf8 | Pbk | Kif5c | Eml5 | Ralyl | Tmem178 |
| Kifc5b | Cttnbp2 | Spdl1 | Chd1l | Rpp30 | Rfx4 | Lhx2 | Trim67 | Adk | Nyap1 |
| Rfc4 | Kif23 | Ints12 | Pak3 | Hopx | Nol4 | Cep72 | Tpd52l2 | Neurod1 | Mpped1 |
| Shcbp1 | 2410004B18Rik | Sdk2 | Kif11 | Ralyl | Rps2 | Epha3 | Lzts1 | Tnik | Syt11 |
| Ckap5 | Ppp3ca | Kif5c | Zmym1 | Gm17750 | Actb | Pak1 | Zdhhc18 | Stxbp1 | Khdrbs2 |
| Apba2 | Hells | Akr7a5 | Lig1 | Ttll1 | Ppfia2 | Gm11266 | Sstr2 | Ripor1 | Tspan12 |
| Sfxn5 | Nol4 | Nrp1 | Hspe1 | Elavl2 | Clstn3 | Josd1 | Dbn1 | Jkamp | Kctd4 |
| Myl12b | B2m | Nav1 | E2f8 | Dtd1 | Fam49a | Ripor1 | Vasp | Taf1 | Adck5 |
| Lzts1 | Chp1 | Nme7 | Afap1l2 | Nr2f1 | Mtif3 | Serpini1 | Entpd1 | Ranbp9 | Rab11b |
| Cdc20 | Dalrd3 | Kif2c | Sptbn1 | Leprotl1 | Trp53inp2 | Snap25 | Kdm5b | Pla2g7 | Cntnap5a |
| Tipin | Nol4l | Elavl4 | Tpx2 | Clybl | Ctnnd2 | Jkamp | Prkab1 | Cplx2 | Rbm15 |
| Nde1 | Siah3 | Nsrp1 | Pclaf | Khdrbs2 | Ttc28 | Kif5a | Clvs1 | Insc | Srrm4 |
| Nsd2 | Myo10 | Aif1l | Ddx19a | S100a10 | Kctd4 | Satb1 | Zfp324 | Rbms1 | Zfpm2 |
| Klf3 | Tox3 | Ptprs | Cenpn | Klf3 | Tspan12 | Jpt1 | Sc5d | Slf1 | Tmem163 |
| Nucks1 | Knl1 | Rfc2 | Cdkn3 | Rfc2 | Cables1 | 2610035D17Rik | Ppfia2 | Gnl2 | Hspe1 |
| Lmnb1 | Tacc3 | H2afv | Cntrob | Ick | Pgm2 | Rcor2 | Palm | Plch1 | Ccdc28b |

| | | | | | | | | | |
|---|---|---|---|---|---|---|---|---|---|
| Cdc25c | Tfap2c | Car11 | Esco2 | Snap25 | Tpd52l2 | Ttyh3 | Tmpo | Tmeff1 | Cnih3 |
| Nxt1 | Nusap1 | Dlgap5 | Nol4 | Cmpk1 | Prmt8 | Slc7a7 | Elmo1 | Brms1l | Satb1 |
| Nusap1 | Khdrbs2 | Smc4 | Slc12a4 | Stxbp1 | Mmd2 | Leprotl1 | Chn2 | Rbfox3 | Dopey1 |
| Cdv3 | Inhbb | Cbs | Hjurp | Nlgn2 | Ralyl | Brms1l | Ttc8 | Dll3 | Rabggtb |
| Dnmt1 | 4930558J18Rik | Tmem238 | Stox1 | Igfbpl1 | Hat1 | 2600014E21Rik | Cttnbp2 | Usp3 | Hecw1 |
| Ube2t | Uhrf1 | Cep72 | Ccdc34 | Usp3 | Rassf3 | Plch1 | Ddah2 | Lims1 | Chp1 |
| Sgo2a | Snap25 | Tmeff1 | Rgma | Idh1 | Csmd2 | Rbfox3 | Cd9 | Sema3c | Glipr2 |
| Rbfox3 | Pitpnc1 | Gm11266 | Trim32 | Diexf | Nrn1 | Sh3kbp1 | Fxyd6 | Kif5a | Id2 |
| Sh3kbp1 | Mipep | Kif15 | Melk | Kmt2a | Spop | Neurod4 | Nav1 | Snhg10 | Csad |
| Hirip3 | Mcm4 | Pon2 | Tuba1c | Rps19 | Pdpn | Rdm1 | Zfp719 | Ahdc1 | Ldha |
| Ptprs | Csmd2 | Itpkb | Mki67 | Ciapin1 | Tubb3 | Syne2 | Dazap1 | Mkrn1 | E2f3 |
| Nedd1 | Thbs3 | Me3 | Top2a | Zfp260 | Hopx | Igsf8 | Galnt17 | Igdcc3 | Fam49a |
| Pmf1 | Pif1 | Nucks1 | Rabggtb | Cenpu | Ppp3ca | Fdxacb1 | Nkain4 | Rasgef1b | Cdk6 |
| Ect2 | Pdia4 | Cttnbp2 | Pdpn | Pak1 | Actn1 | Kdm5b | Usp2 | Pak1 | Lzts1 |
| Cenpc1 | Entpd1 | Frmd5 | Lhx2 | Ctnnd2 | Mxra7 | Kif21b | Myt1l | Slc25a14 | Smc4 |
| Ripk1 | Aftph | Hepacam | Fam161b | Mt2 | Shb | Snhg10 | Bmp1 | Scarb2 | Wasf2 |
| Wee1 | Ago2 | Gabrb1 | Rassf3 | Ipo9 | Lsm6 | Gm17750 | Larp7 | Ptprd | Eml5 |
| Kif23 | Acot13 | Apba2 | Ddah2 | Hnrnpdl | Stmn1 | Myo19 | Gnai2 | Apba2 | Dbn1 |
| Gm40418 | Ctnnd2 | C330027C09Rik | Krtcap2 | Lmnb1 | Prim1 | Ptprd | Dlgap1 | Ciapin1 | Dazap1 |
| Diaph3 | Top2a | Nsg2 | Mgll | Rnf121 | Pccb | Sf1 | Aff2 | Lyrm4 | Gpd1 |
| Mdga1 | Trim32 | Nfib | Zfp608 | Gipc1 | Klhdc3 | Lzts1 | Ppih | Cadm3 | Ncbp2 |
| Mad2l1 | Ston2 | Lix1 | Oat | Atp1b3 | Smc4 | Serinc1 | Wasf2 | Jam3 | Dusp14 |
| Srrm4 | Ccs | Mcm7 | Rab7 | Cep295 | Zfp446 | Lyrm4 | Mdm2 | Nbdy | Actb |
| Cdk5rap2 | Tedc1 | Rasgef1b | Naa50 | Jkamp | Nt5c2 | Pak7 | 2300009A05Rik | Gnai2 | Kif21b |
| Tbc1d31 | Abcf1 | Fam210a | Igfbpl1 | Sox8 | Larp7 | Suclg1 | Abcf1 | Rcor2 | Alcam |
| Arl6ip1 | B3gnt5 | Mt3 | Shroom2 | Tsc22d1 | 2410004B18Rik | Basp1 | Dctn5 | Ick | Nuak1 |
| Atpif1 | Six5 | Ptgds | Eml5 | Scarb2 | Dopey1 | Mtus1 | Neurod4 | Klf3 | Tox3 |
| Cep55 | Lig1 | Mgll | Ston2 | Tnik | Lrrc42 | Gm40418 | Trp53inp2 | Igsf21 | Dpysl3 |
| Parpbp | Atad2 | Gsk3b | Kif23 | Cacybp | Timeless | Zfp91 | Aco1 | Mpped1 | Gap43 |
| Sorbs2 | Rad18 | Bmper | Mtfr2 | Aplp1 | Med4 | Ccdc34 | Retreg3 | Btbd17 | Arf2 |
| Dek | Cit | Fosb | Cdc25c | Golph3l | Dusp10 | Ulk2 | Bcl11a | Sobp | Dhtkd1 |
| Ppwd1 | Lix1 | Dtna | Inhbb | Neo1 | Nsun4 | Taf1 | Foxj3 | Cdk5rap2 | Gm17750 |

Supplemental Table 1
